# ETV4 restrains the formation of CD74^+^ cytotoxic exhausted CD8^+^ T cells

**DOI:** 10.1101/2025.10.02.677509

**Authors:** Dongwook Lee, Yewon Kim, Hyeonmin Gil, Jiyun Lee, Yoon Ha Choi, Minjung Kang, Yunjung Hur, Youngkwon Song, Jong Seok Park, Kwangsoon Kim, Jong Kyoung Kim, Yoontae Lee

**Author notes:** These authors contributed equally (Yewon Kim, Hyeonmin Gil). Corresponding author: Yoontae Lee, Room 388, POSTECH Biotech Center, 77 Cheongam-Ro, Nam-Gu, Pohang, Gyeongbuk 37673, Republic of Korea.

## Abstract

Exhausted CD8^+^ T (Tex) cells comprise heterogeneous subsets with varying cytotoxic and dysfunctional properties, making inhibitory receptor profiles unreliable indicators of function. Therefore, it is essential to define transcriptional regulators or markers that accurately reflect these states. Here, we identified ETS variant 4 (ETV4) as a transcription factor that restricts cytotoxic Tex differentiation within tumors. ETV4 deficiency promotes the emergence of a CD74^+^ cytotoxic Tex subset while preserving the progenitor Tex (Tpex) pool. CD74 serves as both a marker and regulator, reinforcing Tpex features and enhancing Tex cytotoxicity. Mechanistically, ETV4 loss derepresses T-bet expression, thereby increasing IFNγ production that induces CD74 expression in a paracrine manner. Single-cell analyses across human cancers revealed that CD74^+^ Tex cells are enriched for proliferative and metabolic programs and correlate with improved survival. Collectively, we establish the ETV4–CD74 axis as a regulator of Tex fate and promising therapeutic target to augment anti-tumor immunity.

## Introduction

CD8^+^ T cells are key effectors for eliminating tumors and virus-infected cells. However, under conditions of chronic antigen exposure, CD8^+^ T cells progressively lose their ability to secrete effector cytokines such as IL-2, IFNγ, and TNF, exhibit reduced proliferative capacity, and undergo increased apoptosis. This overall loss of effectiveness and population is termed exhaustion^1, 2, 3^. Within this program, TCF-1^+^SLAMF6^+^ progenitor exhausted T (Tpex) cells sustain long-term responses through self-renewal. Persistent antigenic stimulation drives Tpex cells to gradually differentiate into PD-1^+^TIM3^+^ exhausted T (Tex) cells through successive rounds of proliferation^4, 5^. Compared with Tpex, Tex cells express higher levels of multiple inhibitory receptors (IRs) and display impaired functionality^6^. However, despite this dysfunction, Tex cells retain granzyme B expression and serve as the primary effectors mediating tumor elimination^7^. Immune checkpoint inhibitors, such as anti-PD-1, exploit this property by enhancing the cytotoxicity of CD8^+^ T cells by promoting the differentiation of Tpex cells to Tex cells^4, 8, 9^.

Numerous studies have attempted to delineate the mechanisms underlying CD8^+^ T cell exhaustion by investigating the transcription factors, including TOX, Blimp1, T-bet, Eomes, and BATF, that orchestrate this process^10, 11, 12, 13, 14, 15^. Although TOX is closely associated with the dysfunctional phenotype, its deletion is not an effective strategy to prevent functional exhaustion as it disrupts the establishment of the Tpex population and compromises Tex cell survival^10, 16^. Likewise, although PD-1 is a central IR that restrains T-cell functionality, complete ablation of PD-1 expression leads to immunopathology that outweighs potential therapeutic benefits. In contrast, selective disruption of the −23.8 kb *Pdcd1* enhancer, which is critical for sustaining high PD-1 expression levels in Tex cells, maintains immune regulation while enhancing anti-tumor immunity^17, 18^. These findings underscore the importance of identifying regulators specifically acting at the terminal stage of exhaustion, which may provide more precise therapeutic strategies. ETV4, a well-established oncogenic transcription factor in various cancers^19, 20, 21, 22, 23^, has recently been implicated in T cell exhaustion. Elevated ETV4 activity has been detected in tumor-infiltrating Tex cells, with its DNA-binding motif highly enriched in the dysfunctional Tex population^24, 25, 26^. Furthermore, ETV4 expression is negatively correlated with cytotoxic markers^24^, suggesting a potential role in regulating terminal T cell exhaustion.

Restoring Tex cell functionality is of critical importance; however, accurate evaluation requires surface markers that are reliable indicators of the state of Tex cells, as reliance on IR expression alone can be misleading^27^. To account for Tex heterogeneity, subsets have been delineated by markers, such as CX3CR1^+^ (Tex-int) and CX3CR1^+^KLRG1^+^ (Tex-KLR, a functionally superior subset), whereas CD38 and CD101 identify terminally exhausted CD8⁺ T (Tex-term) cells^28, 29^. However, these markers do not consistently reflect the functional states of Tex cells across single-cell RNA sequencing (scRNA-seq) datasets or in tumor models such as multiple myeloma and melanoma^30^. Therefore, additional markers that accurately define Tex cell functionality remain to be identified.

In this study, we identified ETV4 as a key transcription factor that regulates the differentiation of cytotoxic Tex subsets. Loss of ETV4 was found to lead to increased IR expression; however, it also simultaneously enhanced the polyfunctional cytotoxicity of Tex cells. In addition, ETV4 deficiency elevated CD74 expression in both tumor-infiltrating Tpex and Tex cells. We further demonstrate that CD74 serves as a functional marker of these subsets. Collectively, our findings highlight the ETV4–CD74 axis as a potential therapeutic target to enhance anti-tumor immunity.

## Results

### *Etv4* is predominantly expressed and activated in tumor-infiltrating *Pdcd1*^hi^*Havcr2*^+^ CD8^+^ T cells

Given the potential role of ETV4 in T cell exhaustion^24, 25, 26^, we examined its expression profile using publicly available scRNA-seq data of blood-circulating and tumor-infiltrating CD44^+^ effector CD8^+^ T cells from the MC38 colon carcinoma model^31^ to determine its subset specificity. *Etv4* expression was enriched in tumor-infiltrating *Pdcd1*^hi^*Havcr2*^+^ Tex clusters (clusters 2, 4, 6, 7, and 9) compared with *Pdcd1*^+^*Tcf7*^+^ Tpex cells (clusters 3 and 5) and blood-circulating CD8^+^ T cell clusters (Fig. 1a-c). Similarly, ETV4 transcriptional activity was elevated in tumor-infiltrating *Pdcd1*^hi^*Havcr2*^+^ Tex clusters (Fig. 1d). Notably, however, *Etv4* expression and transcriptional activity were not uniformly distributed across all *Pdcd1*^hi^*Havcr2*^+^ clusters (Fig. 1c, d), implying a subset-specific role of ETV4 within the heterogeneous Tex cell population.

**Figure 1.**
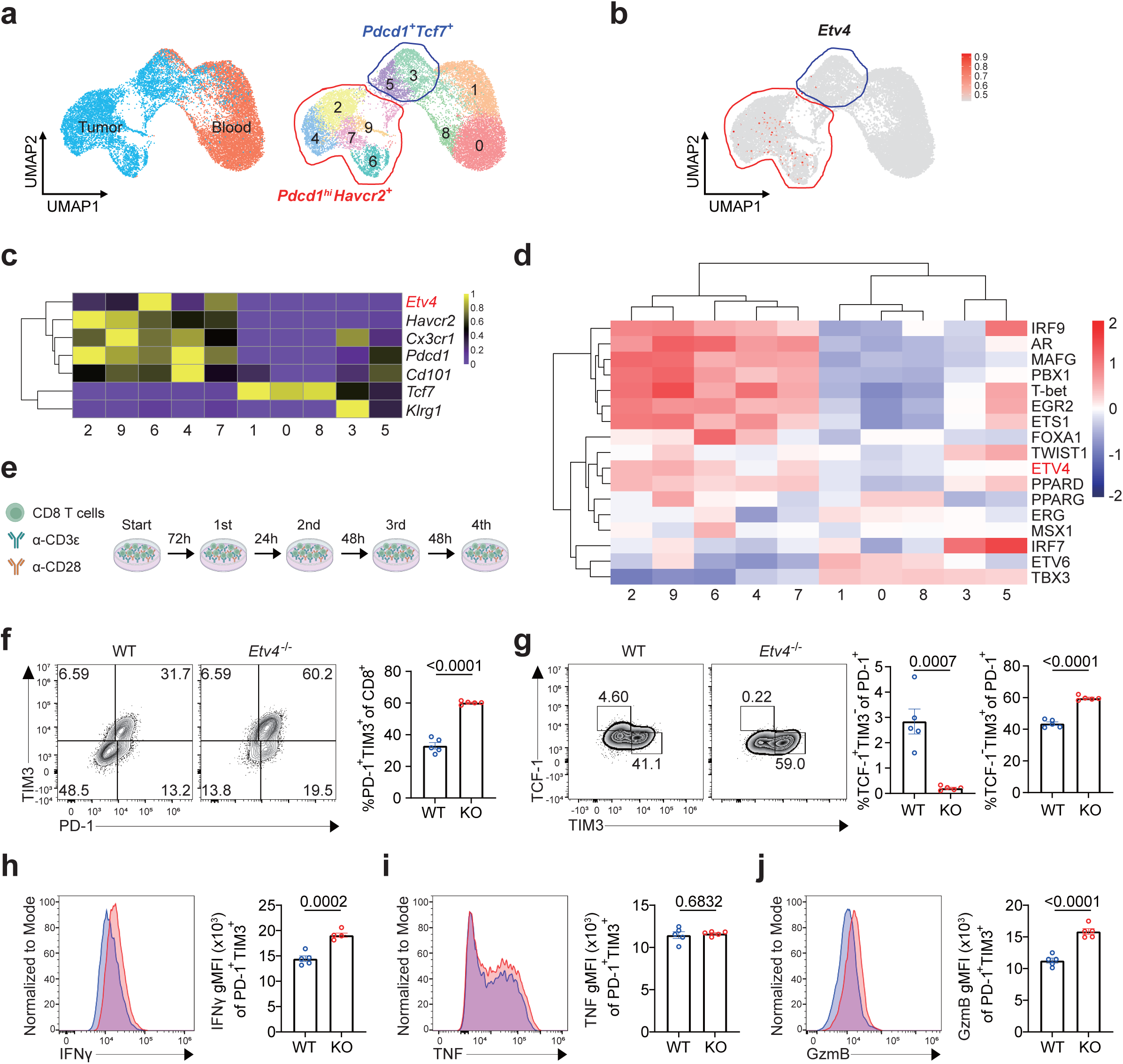
*Etv4* is mainly expressed and activated in *Pdcd1*^hi^ *Havcr2*^+^ CD8^+^ T cells. **a**, uniform manifold approximation and projection (UMAP) clustering of scRNA-seq data from tumor-infiltrating and blood-circulating CD44^+^ effector CD8^+^ T cells in the MC38 colon carcinoma model^31^. **b-d**, Visualization of gene expression and transcription factor activity: feature plot of *Etv4* expression (**b**), heatmap of exhaustion marker expression (**c**), and heatmap of transcription factor activity (**d**). **e**, Schematic of the *in vitro* T cell exhaustion models. **f**, Representative flow cytometry plots and quantification of WT (n=5) and *Etv4* KO (n=5) PD-1^hi^TIM3^+^ CD8^+^ T cells after the fourth round of *in vitro* T cell exhaustion models. **g**, Representative flow cytometry plots and quantification of Tpex (TCF-1^+^TIM3^-^) and Tex (TCF-1^-^TIM3^+^) subsets in WT (n=5) and *Etv4* KO (n=5) cells after the fourth round of *in vitro* T cell exhaustion models. **h-j**, Representative histograms and quantification of geometric mean fluorescence intensity (gMFI) of IFNγ (**h**), TNF (**i**), and granzyme B (**j**) in WT (n=5) and *Etv4* KO (n=5) PD-1^hi^TIM3^+^ CD8^+^ T cells after the fourth round of *in vitro* T cell exhaustion models, restimulated with phorbol 12-myristate 13-acetate (PMA) and ionomycin. Data in **f-j** are representative of three independent experiments. Error bars indicate mean ± s.e.m. Statistical analysis was performed using unpaired two-tailed Student’s t-tests for **f-j**. Panel **e** was created using <u>BioRender.com</u>

To assess the role of ETV4 in Tex differentiation, we subjected wild-type (WT) and *Etv4* knockout (KO) CD8^+^ T cells to consecutive rounds of chronic T cell receptor (TCR) stimulation *in vitro*, mimicking T cell exhaustion^12, 32^ (Fig. 1e and Extended Data Fig. 1a, b). ETV4 expression was induced in CD8^+^ T cells following chronic TCR stimulation but was absent in *Etv4* KO CD8^+^ T cells (Extended Data Fig. 1c). ETV4 loss increased the frequency of PD-1^+^TIM3^+^ Tex cells while decreasing that of TIM3^-^TCF-1^+^ Tpex cells (Fig. 1f, g). Despite the enhanced exhaustion phenotype, *Etv4* KO PD-1^+^TIM3^+^ Tex cells exhibited greater cytotoxicity than their WT counterparts (Fig. 1h-j).

Tpex cells sustain the overall Tpex–Tex pool through self-renewal and differentiation into Tex cells^4, 33^. As *Etv4*-deficient CD8^+^ T cells exhibited a reduced Tpex frequency (Fig. 1g), we examined their ability to maintain population size under chronic TCR stimulation. In co-culture experiments using congenically marked WT and *Etv4* KO CD8^+^ T cells under identical stimulation conditions (Extended Data Fig. 1d), both populations maintained their initial frequencies through four rounds of TCR stimulation, indicating that *Etv4*-deficient cells are competitively fit and not outcompeted by their WT counterparts (Extended Data Fig. 1e). Moreover, *Etv4* KO CD8^+^ T cells exhibited increased cytotoxic activity and greater resistance to apoptosis than WT CD8^+^ T cells (Extended Data Fig. 1f, g), demonstrating enhanced resilience to functional exhaustion. Together, these findings reveal an interesting paradox wherein ETV4 loss promotes the differentiation of CD8^+^ T cells toward Tex cells, but simultaneously protects them from functional exhaustion by reducing apoptosis and reinforcing cytotoxicity.

### Loss of ETV4 enhances CD8^+^ T cell functionality in tumors

Given the regulation of CD8^+^ T cell exhaustion and functionality by ETV4 *in vitro*, we next investigated its role in tumor-infiltrating CD8^+^ T cells *in vivo*. We performed scRNA-seq on CD8^+^ T cells isolated from MC38 tumors in WT and *Etv4* KO mice and identified major PD-1^+^ CD8^+^ T cell subsets, including Tpex, Tex-ISG (IFN-stimulated genes), Tex-pro (proliferating), Tex-int, Tex-KLR, and Tex-term (Fig. 2a and Extended Data Fig. 2a-f). The relative composition of these subsets differed significantly between WT and *Etv4* KO mice (Fig. 2b, c). In particular, *Etv4* KO mice exhibited an expansion of Tex-KLR cells accompanied by a reduction of Tex-int cells, whereas the frequency of Tpex cells was largely comparable with that in WT mice (Fig. 2c). In naïve *Etv4* KO mice, the splenic immune compartment—including natural killer (NK) cells, B cells, CD4^+^ and CD8^+^ T cells, regulatory CD4^+^ T cells, macrophages, conventional dendritic cells (DCs), monocytes, and neutrophils—was unaffected in comparison with that in WT mice (Extended Data Fig. 3a-h), indicating that ETV4 is dispensable for steady-state immune homeostasis. Further, the overall composition of tumor-infiltrating immune cells, proliferative capacity of CD4^+^ and CD8^+^ T cells, and cytotoxic activity of NK and CD4^+^ cells were unchanged in *Etv4* KO mice (Extended Data Fig. 4a-m). However, despite all these similarities in immune cell composition between *Etv4* KO and WT mice, *Etv4* KO mice displayed superior tumor growth control (Fig. 2d, e).

**Figure 2.**
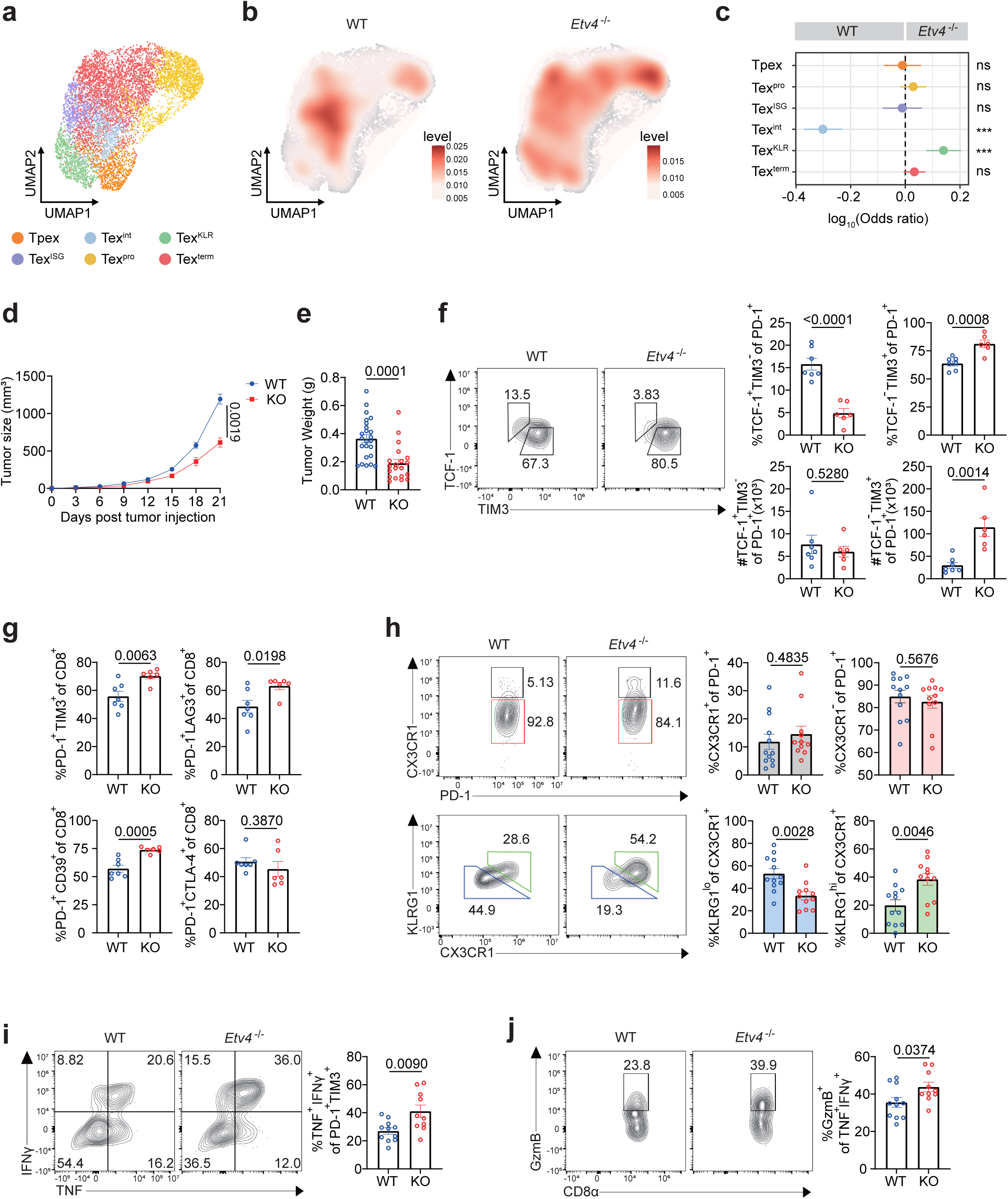
Loss of ETV4 enhances CD8^+^ T cell functionality. **a,** UMAP dimensional reduction of scRNA-seq data from MC38 tumor-infiltrating WT and *Etv4* KO CD8^+^ T cells. **b,** UMAP plots showing cluster density of WT versus *Etv4* KO CD8⁺ T cells. **c,** Bar graph of cluster proportions in WT and *Etv4* KO CD8⁺ T cells. **d,** Tumor growth curves in MC38 tumor-bearing WT (n=5) and *Etv4* KO (n=5) mice. **e,** Tumor weight of MC38 tumor-bearing WT (n=24) and *Etv4* KO (n=20) mice. **f,** Representative flow cytometry plots and quantification of Tpex (TCF-1^+^TIM3^-^) or Tex (TCF-1^-^TIM3^+^) cells infiltrating MC38 tumors in WT (n=7) and *Etv4* KO (n=6) mice. **g,** Flow cytometric quantification of PD-1^hi^TIM3^+^, PD-1^hi^CD39^+^, PD-1^hi^LAG3^+^, and PD-1^hi^CTLA-4^+^ CD8^+^ T cells infiltrating MC38 tumors in WT (n=7) and *Etv4* KO (n=6) mice. **h,** Representative flow cytometry plots and quantification of PD-1^+^CX3CR1^+^, PD-1^+^CX3CR1^+^ KLRG1^-^ (Tex-int), PD-1^+^CX3CR1^+^KLRG1^+^ (Tex-KLR), and PD-1^+^CX3CR1^-^ (Tex-term) CD8^+^ T cells infiltrating MC38 tumors in WT (n=12) and *Etv4* KO (n=11) mice. **i,** Representative flow cytometry plots and quantification of TNF^+^IFNγ^+^ frequencies within PD-1^hi^TIM3^+^ CD8^+^ T cell populations infiltrating MC38 tumors in WT (n=11) and *Etv4* KO (n=10) mice, restimulated with PMA and ionomycin. **j,** Representative flow cytometry plots and quantification of granzyme B^+^ frequencies within TNF^+^IFNγ^+^PD-1^hi^TIM3^+^ CD8^+^ T cells infiltrating MC38 tumors in WT (n=11) and *Etv4* KO (n=10) mice, restimulated with PMA and ionomycin. Data represent five independent experiments for **d** and three for **f-j**. Data were pooled from five independent experiments for **e** and from two independent experiments for **h-j**. Error bars indicate mean ± s.e.m. Statistical significance was determined using a two-way ANOVA with Tukey’s multiple comparison for **d** and unpaired two-tailed Student’s t-tests for **e-j**.

Within the CD8^+^ T cell compartment, ETV4 deficiency skewed differentiation toward the TIM3^+^TCF-1^-^ Tex phenotype, accompanied by a reduced frequency of TIM3^-^TCF-1^+^ Tpex cells (Fig. 2f), consistent with our *in vitro* observations (Fig. 1g). Notably, in tumors of *Etv4* KO mice, the absolute number of Tpex cells was preserved, whereas the number of Tex cells was significantly increased compared with those in tumors of WT mice (Fig. 2f). *Etv4* KO CD8^+^ T cells also showed broad expansion of IR-expressing subsets, with the exception of CTLA-4 (Fig. 2g). In line with the scRNA-seq results (Fig. 2c), ETV4 deficiency preferentially promoted differentiation toward the cytotoxic Tex-KLR subset while reducing Tex-int populations, without changes in Tex-term populations (Fig. 2h). As the overall frequency of CX3CR1^+^ cells did not differ between *Etv4* KO and WT mice, these results indicate a compositional shift within this population from Tex-int to Tex-KLR cells. Reflecting the cytotoxic property of Tex-KLR^29^, *Etv4* KO PD-1^+^TIM3^+^ Tex cells exhibited an enhanced cytotoxic profile (Fig. 2i, j), mirroring the phenotype observed *in vitro* (Fig. 1h, j).

To confirm that these effects were T cell intrinsic, we performed adoptive transfer experiments using ovalbumin (OVA)-specific OT-I CD8^+^ T cells. Intratumoral transfer of pre-activated *Etv4* KO OT-I cells resulted in superior tumor control than that of WT OT-I cells (Extended Data Fig. 5a, b), demonstrating that the functional advantage conferred by *Etv4* loss was intrinsic to CD8^+^ T cells. To further assess the intrinsic role of ETV4 in regulating CD8^+^ T cell functionality within the same tumor microenvironment (TME), we co-adoptively transferred congenically marked WT and *Etv4* KO OT-I cells into *Rag1* KO mice bearing B16-OVA tumors. In this setting, *Etv4* KO OT-I cells exhibited increased proliferation, reduced apoptosis, and enhanced cytotoxicity (Extended Data Fig. 5c-e). Collectively, these results demonstrate that *Etv4*-deficient CD8^+^ T cells resist multiple hallmarks of functional exhaustion while differentiating into Tex cells, establishing ETV4 as a negative regulator of CD8^+^ T cell functionality in tumors.

### ETV4 regulates the differentiation of CD74-expressing exhausted CD8^+^ T cells

As *Etv4* KO Tex cells displayed both elevated IR expression and enhanced cytotoxicity, we reasoned that IR profiles alone may be insufficient to explain functional heterogeneity. Therefore, we sought to identify surface markers regulated by ETV4 that could better predict Tex functionality. To improve the resolution of ETV4 activity within heterogenous tumor-infiltrating CD8^+^ T cells and thereby define ETV4-regulated genes with greater reliability, the initial 6 clusters from our scRNA-seq dataset were further subdivided into 10 clusters (Fig. 3a). Among these, ETV4 activity was the highest in proliferating Tex cluster 1 (Fig. 3a), corresponding to the proliferative burst during the Tpex-to-Tex transition^4, 5^. Gene set enrichment analysis (GSEA) of cluster 1 revealed significant enrichment of IFNγ response genes, including *Cd74*, in *Etv4* KO CD8^+^ T cells (Fig. 3b), consistent with our observations that *Etv4*-deficient CD8^+^ T cells produce elevated IFNγ both *in vitro* and *in vivo* (Fig. 1h and 2i).

**Figure 3.**
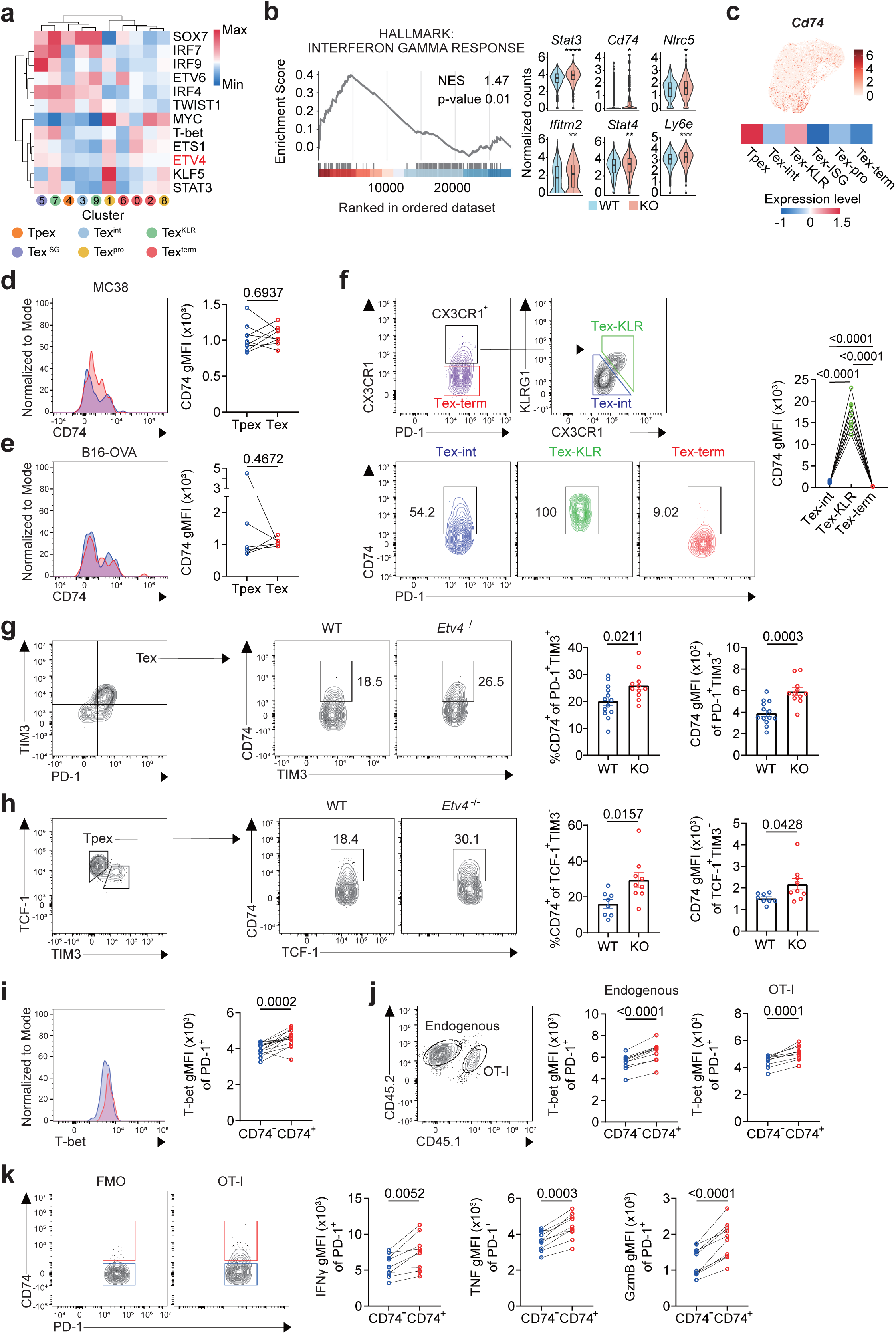
ETV4 regulates the differentiation of CD74-expressing exhausted CD8^+^ T cells. **a,** Heatmap of transcription factor activity from scRNA-seq of MC38 tumor-infiltrating WT versus *Etv4* KO CD8*^+^* T cells. **b,** Gene set enrichment analysis (GSEA) of selected immunological signature and hallmark gene sets in *Etv4* KO Tex cluster 1 compared with WT Tex cluster 1. **c,** *Cd74* expression (red) in MC38-infiltrating WT CD8*^+^* T cells, shown by feature plot and heatmap. **d-f,** Representative histograms, flow cytometry plots, and gMFI quantification of CD74 surface expression in tumor-infiltrating CD8^+^ T cells: Tpex (TCF-1^+^TIM3^-^) versus Tex (TCF-1^-^TIM3^+^) subsets from MC38 (n=9) (**d**) and B16-OVA (n=6) (**e**) tumors, and Tex subsets defined as PD-1^+^CX3CR1^+^KLRG1^-^ (Tex-int), PD-1^+^CX3CR1^+^KLRG1^+^ (Tex-KLR), and PD-1^+^CX3CR1^-^ (Tex-term) from MC38 tumors (n=9) (**f**). **g,** Representative flow cytometry plots and percentage/gMFI quantification of CD74 surface expression in PD-1^hi^TIM3^+^ Tex cells infiltrating MC38 tumors in WT (n=13) and *Etv4* KO (n=11) mice. **h,** Representative flow cytometry plots and percentage/gMFI quantification of CD74 surface expression in TCF-1^+^TIM3^−^ Tpex cells from B16-OVA tumor-bearing *Rag1*KO mice intratumorally adoptively transferred with WT OT-I (n=8) or *Etv4* KO OT-I (n=9) cells. **i,** Representative histograms and gMFI quantification of T-bet intracellular expression in CD74^−^ (blue) versus CD74^+^ (red) PD-1^+^ CD8^+^ T cells from MC38 tumors (n=16). **j,** Representative flow cytometry plots and gMFI quantification of intracellular T-bet expression in CD74^−^ versus CD74⁺ PD-1⁺ T cells from CD45.2 endogenous CD8^+^ T and intratumorally adoptively transferred CD45.1/2 OT-I cells in B16-OVA tumors of CD45.2 immunocompetent mice (n=10). **k,** Representative flow cytometry plots and gMFI quantification of intracellular IFNγ, TNF, and granzyme B expression in CD74⁻ (blue) versus CD74^+^ (red) CD45.1/2 PD-1^+^ OT-I cells intratumorally adoptively transferred into B16-OVA tumors of CD45.2 immunocompetent mice (n=10) after restimulation with OVA 257-264 peptide. Data represent three independent experiments for **d-k**. Error bars indicate mean ± s.e.m. Statistical significance was determined using paired two-tailed Student’s t-tests for **d-f** and **i-k,** and unpaired two-tailed Student’s t-tests for **g-h.** Data were pooled from two independent experiments for **f-i**.

CD74, originally described as an invariant chain involved in antigen presentation^34^, also interacts with macrophage migration inhibitory factor (MIF), which has been reported to inhibit CD8^+^ T cell proliferation and granzyme B production^35^. Although CD74 is highly expressed in macrophages, DCs, and B cells, emerging evidence shows that it is also expressed in regulatory CD4^+^ and CD8^+^ T cells within the TME^36, 37, 38, 39^. The paradoxical coexistence of enhanced cytotoxicity and elevated *Cd74* expression in *Etv4* KO CD8^+^ T cells prompted us to further investigate the role of CD74 in Tex cells. Tumor-infiltrating CD8⁺ T cells expressed CD74 at higher levels than those from spleen or tumor-draining lymph nodes (Extended Data Fig. 6a). In scRNA-seq data, *Cd74* expression was confined to Tpex and Tex-KLR subsets (Fig. 3c). Although no major difference was detected between TIM3^-^TCF-1^+^ Tpex and TIM3^+^TCF-1^-^ Tex subsets in both MC38 and B16-OVA tumors (Fig. 3d, e), CD74 expression was enriched in Tex-KLR cells within the heterogeneous Tex population of MC38 tumors (Fig. 3f). Aligning with the CD74 expression pattern observed in MC38 tumor-infiltrating CD8^+^ T cells (Fig. 3f), *in vitro* chronic TCR stimulation induced the highest CD74 expression during the second and third cycles, followed by a sharp decline at the fourth (Extended Data Fig. 6b). Furthermore, CD74 expression was higher in *Etv4* KO CD8^+^ T cells than in WT cells, at both protein and mRNA levels under *in vitro* chronic TCR stimulation (Extended Data Fig. 6c, d). This upregulation was also observed in *Etv4* KO PD-1^+^TIM3^+^ Tex cells from MC38 tumors (Fig. 3g) and in *Etv4* KO TCF-1^+^TIM3^-^ Tpex cells from B16-OVA tumors (Fig. 3h), indicating that ETV4 deficiency influences CD74 expression in both Tpex and Tex compartments.

Next, we examined transcriptional regulators associated with T cell exhaustion in CD74^+^ T cells. We directly compared CD74^+^ and CD74^−^ T cells and found that, among T-bet, TOX, and Eomes, only T-bet was significantly increased in PD-1^+^CD74^+^ CD8^+^ T cells infiltrating MC38 tumors, as well as in PD-1^+^CD74^+^ endogenous CD8^+^ T and OT-I cells infiltrating B16-OVA tumors, compared with their CD74^−^ counterparts (Fig. 3i, j and Extended Data Fig. 6e-h). T-bet sustains Tex effector function and limits their conversion into terminally exhausted resident CD8⁺ T cells^13, 40^. The elevated T-bet expression in CD74^+^ T cells was reflected by the enhanced cytotoxicity of B16-OVA tumor-infiltrating CD74^+^PD-1^+^ OT-I cells compared with their CD74^−^ counterparts (Fig. 3k). This observation aligns with earlier findings in the context of viral infection, where CD74^+^ CD8^+^ T cells also exhibited increased granzyme B levels^41^, supporting CD74 as a conserved functional marker across disease contexts. Collectively, these findings establish CD74 as a robust marker that reflects Tex heterogeneity and as a potential prognostic biomarker under ETV4-mediated regulation.

### CD74 preserves Tpex pools and cytotoxicity

Next, we investigated whether CD74 serves solely as a surface marker or also contributes to Tex cell function. To this end, we co-cultured congenically marked CD8^+^ T cells transduced with either *Cd74*-overexpressing or empty vector retrovirus in the absence or presence of MIF, a well-characterized ligand for CD74 (Fig. 4a and Extended Data Fig. 7a). Unexpectedly, CD74 overexpression selectively expanded the TIM3⁻TCF-1^+^ Tpex population without affecting TIM3^+^TCF-1⁻ Tex cells (Fig. 4b) and also enhanced cytotoxicity (Fig. 4c, d), indicating that CD74 plays an active role in sustaining Tpex cells and promoting effector function. However, MIF treatment failed to augment these effects (Fig. 4b-d), suggesting that the CD74-mediated regulation of Tpex cells and cytotoxicity is independent of MIF engagement.

**Figure 4.**
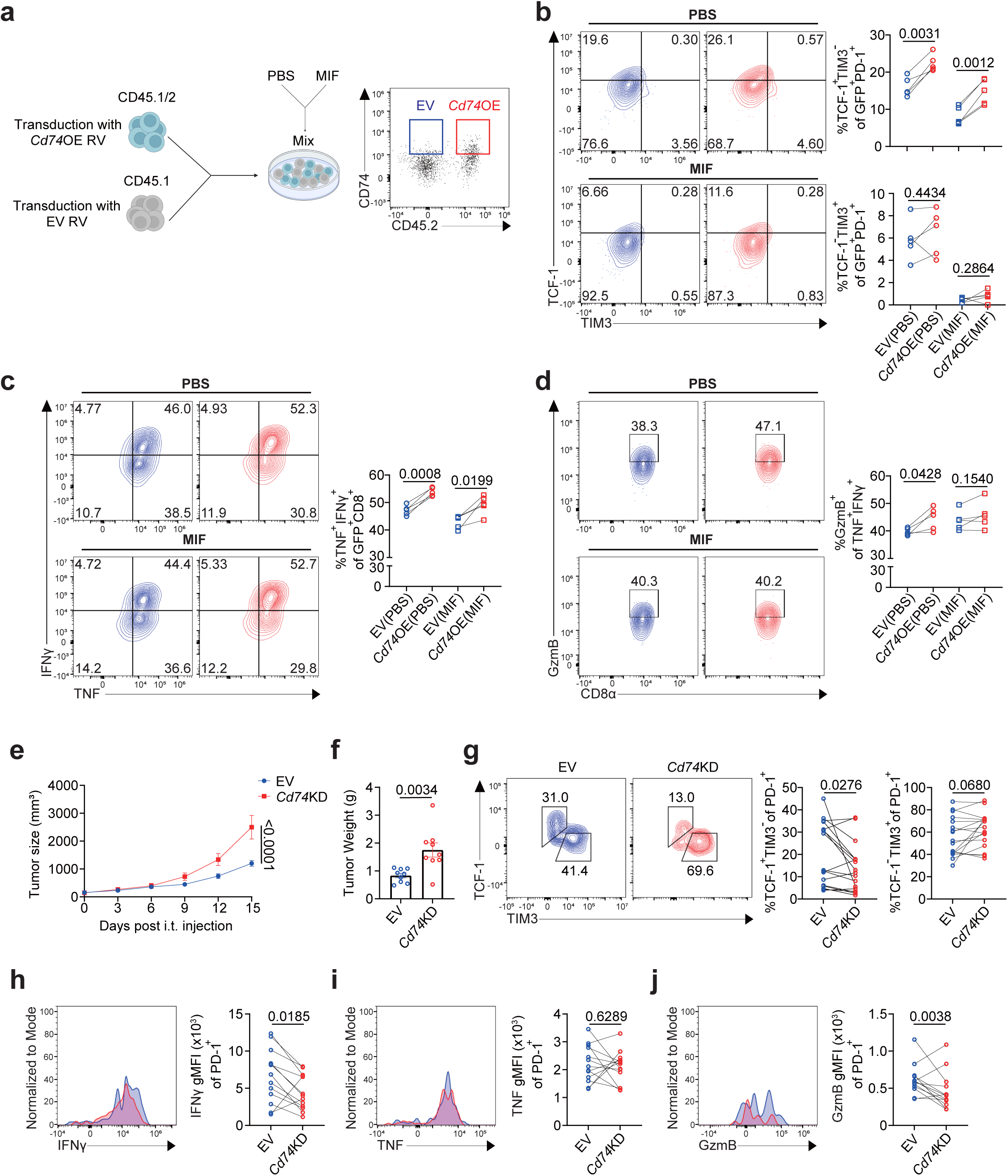
CD74 preserves Tpex pools and cytotoxicity. **a,** Experimental design of co-culture assays in which CD45.1 empty vector (EV) and CD45.1/2 *Cd74-*overexpressing (OE) CD8^+^ T cells underwent two rounds of continuous TCR stimulation (one before and one after transduction, with the latter treated with PBS or MIF), and representative flow cytometry plots of CD45.1 EV (blue) and CD45.1/2 *Cd74* OE (red) CD8^+^ T cells. **b,** Representative flow cytometry plots and quantification of Tpex (TCF-1^+^TIM3^-^) and Tex (TCF-1^-^TIM3^+^) subsets from co-cultured CD45.1 EV (blue) and CD45.1/2 *Cd74* OE (red) CD8⁺ T cells after two rounds of continuous TCR stimulation (one before and one after transduction, with the latter treated with PBS or MIF) (n=5). **c-d**, Representative flow cytometry plots and quantification of CD8^+^ T cells following two rounds of continuous TCR stimulation (one before and one after transduction, with the latter treated with PBS or MIF), restimulated with PMA and ionomycin (n=5): TNF^+^IFNγ^+^ cell frequencies (**c**), and granzyme B^+^ frequency of TNF^+^IFNγ^+^ cells (**d**). **e-f,** Tumor growth curves (**e**) and tumor weight (**f**) in B16-OVA tumor-bearing immunocompetent mice intratumorally adoptively transferred with EV OT-I (n=9) or *Cd74* KD OT-I (n=9) cells. **g,** Representative flow cytometry plots and quantification of Tpex (TCF-1^+^TIM3^-^) or Tex (TCF-1^-^ TIM3^+^) subsets in B16-OVA tumor-bearing CD45.1 immunocompetent mice after intratumoral co-adoptive transfer of CD45.2 EV (blue) and CD45.1/2 *Cd74* knockdown (KD) OT-I T cells (red) (n=17). **h-j**, Representative histograms and gMFI quantification of PD-1^+^ OT-I cells in B16-OVA tumor-bearing CD45.1 immunocompetent mice after intratumoral co-adoptive transfer of CD45.2 EV (blue) and CD45.1/2 *Cd74* KD OT-I T cells (red) (n=13), restimulated with OVA 257-264 peptide: IFNγ **(h)**, TNF **(i)**, and granzyme B **(j)**. Data represent three independent experiments for **b-j**. Error bars indicate mean ± s.e.m. Statistical significance was determined using paired two-tailed Student’s t-tests for **b-d** and **g-j,** unpaired two-tailed Student’s t-tests for **f**, and two-way ANOVA with Tukey’s multiple comparison for **e.** Data in **g** were pooled from two independent experiments. Panel **a** was created using <u>BioRender.com</u>

To assess the role of endogenous CD74, we adoptively transferred pre-activated OT-I cells transduced with the *Cd74*-knockdown (*Cd74* KD) or empty vector into B16-OVA tumor-bearing immunocompetent mice (Extended Data Fig. 7a, b). Compared with controls, *Cd74* KD OT-I cells exhibited impaired tumor control (Fig. 4e, f). In competitive co-transfer experiments, *Cd74* KD OT-I cells showed a marked reduction in TIM3^-^TCF-1^+^ Tpex formation (Fig. 4g), consistent with the results of CD74 overexpression *in vitro* (Fig. 4b). As CD74 expression is initiated within Tpex cells (Fig. 3c), these results identify CD74 as a gatekeeper for maintaining Tpex populations. This mechanism may also explain why *Etv4*-deficient CD8^+^ T cells, which display increased CD74 expression, preserve Tpex pools despite expanded Tex populations (Fig. 2c, f).

Furthermore, *Cd74* KD OT-I cells expressed lower levels of IFNγ and granzyme B than controls (Fig.4h-j), whereas proliferation and apoptosis were comparable with those of controls (Extended Data Fig. 7c, d). Collectively, these results demonstrate that CD74 is not a passive marker but an active regulator of exhaustion differentiation that preserves the Tpex pool and reinforces cytotoxic function to support durable anti-tumor responses.

### ETV4 deficiency promotes CD74 upregulation through increased IFNγ production

To elucidate the mechanism underlying CD74 upregulation in *Etv4*-deficient CD8^+^ T cells, we investigated whether this is a cell-intrinsic effect or mediated by extrinsic cues. We employed two competitive experimental systems: (1) co-culture of congenically marked WT and *Etv4* KO CD8^+^ T cells under chronic *in vitro* TCR stimulation; and (2) co-adoptive transfer of congenically marked naïve WT and *Etv4* KO OT-I cells into B16-OVA tumors. In both settings, CD74 expression levels were comparable between WT and *Etv4* KO cells (Fig. 5a, b), suggesting regulation by extrinsic factors. We next examined whether *Etv4* KO CD8^+^ T cells could induce CD74 expression in neighboring WT CD8^+^ T cells. For this, CD45.1/2 WT CD8^+^ T cells (responders) were co-cultured or co-transferred with either CD45.2 WT or CD45.2 *Etv4* KO CD8^+^ T cells (Fig. 5c). Strikingly, *Etv4* KO CD8^+^ T cells promoted the emergence of CD74^+^ WT bystanders (Fig. 5d-e), suggesting that paracrine signaling contributes to enhanced CD74 expression.

**Figure 5.**
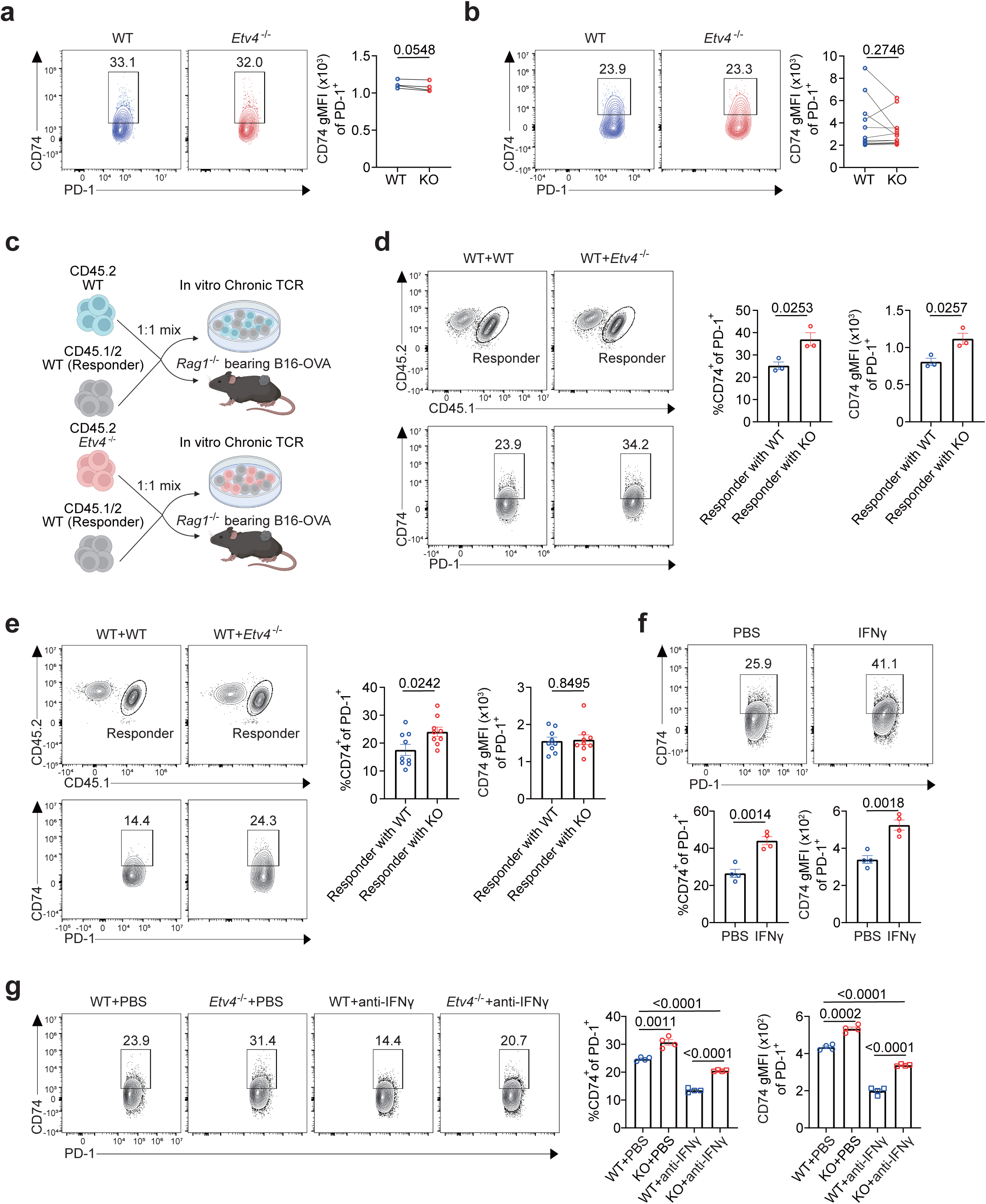
ETV4 deficiency promotes CD74 upregulation through increased IFNγ production. **a,** Representative flow cytometry plots and gMFI quantification of CD74 surface expression in PD-1^+^ CD8^+^ T cells at the third activation cycle during the *in vitro* T cell exhaustion models in co-cultures of CD45.1/2 WT (blue) and CD45.2 *Etv4* KO (red) CD8^+^ T cells (n=4). **b,** Representative flow cytometry plots and gMFI quantification of CD74 surface expression in PD-1^+^ CD8^+^ T cells from B16-OVA tumor-bearing *Rag1* KO mice after intratumoral co-adoptive transfer of CD45.1 WT (blue) and CD45.2 *Etv4* KO OT-I cells (red) (n=12). **c,** Experimental outline for *in vitro* co-culture of CD45.1 WT CD8^+^ responder cells with either CD45.2 WT or CD45.2 *Etv4* KO CD8^+^ T cells in the T cell exhaustion models (**d**), and intratumoral co-adoptive transfer of CD45.1 WT OT-I responder cells with either CD45.2 WT or CD45.2 *Etv4* KO OT-I cells into B16-OVA tumor-bearing *Rag1* KO mice (**e**). **d-e,** Representative flow cytometry plots and percentage/gMFI quantification of CD74 surface expression in PD-1^+^ CD8^+^ T cells after the third activation in the *in vitro* T cell exhaustion models (n=3) **(d)**, and from B16-OVA tumor-bearing *Rag1* KO mice (n=9) **(e)**. **f,** Representative flow cytometry plots and percentage/gMFI quantification of CD74 surface expression in PD-1^+^ CD8^+^ T cells at the third activation under PBS or IFNγ treatment in the *in vitro* T cell exhaustion models (n=4). **g,** Representative flow cytometry plots and percentage/gMFI quantification of CD74 surface expression in PD-1^+^ CD8^+^ T cells at the third activation of *in vitro* T cell exhaustion models in WT (n=4) and *Etv4* KO (n=4) cells under PBS or anti-IFNγ treatment. Data represent three independent experiments for **a-b** and **d-g**. Error bars indicate mean ± s.e.m. Statistical significance was determined using paired two-tailed Student’s t-tests for **a-b** and unpaired two-tailed Student’s t-tests for **d-g.** Data in **e** were pooled from two independent experiments. Panel **c** was created using <u>BioRender.com</u>

IFNγ has been reported to induce CD74 expression in melanoma cells^42^; therefore, we tested whether it plays a similar role in CD8^+^ T cells. Indeed, we found that IFNγ treatment increased CD74 expression under chronic TCR stimulation (Fig. 5f). Given that *Etv4* KO CD8^+^ T cells produced elevated levels of IFNγ both *in vitro* and *in vivo* (Fig.1h and 2i), we hypothesized that increased IFNγ secretion drives CD74 upregulation in *Etv4* KO CD8^+^ T cells. To test this, we treated WT and *Etv4* KO CD8^+^ T cells undergoing chronic TCR stimulation with an anti-IFNγ neutralizing antibody at the third activation cycle, when differences in CD74 expression were most pronounced (Extended Data Fig. 6c, d). IFNγ neutralization reduced CD74 expression in *Etv4* KO CD8^+^ T cells to levels below those in WT cells (Fig. 5g). Taken together, these findings demonstrate that increased IFNγ production by *Etv4*-deficient CD8^+^ T cells promotes CD74 upregulation through paracrine signaling, underscoring the critical role of extrinsic cytokine cues in shaping T cell phenotypes within tumors.

### ETV4 suppresses T-bet expression in Tex cells

We next sought to identify the molecular basis for the elevated IFNγ expression in tumor-infiltrating *Etv4*-deficient CD8^+^ T cells. Differential expression analysis of our scRNA-seq dataset revealed that *Tbx21* transcript levels were significantly upregulated in *Etv4* KO CD8^+^ T cells (Fig. 6a). *Tbx21* encodes T-bet, a key transcription factor that sustains effector function in Tex cells and promotes IFNγ production in both CD4^+^ and CD8^+^ T cells^13, 40, 43, 44, 45, 46, 47, 48, 49^. Consistent with the change in *Tbx21* levels, T-bet protein expression was also elevated in MC38 tumor-infiltrating *Etv4* KO CD8^+^ T cells (Fig. 6b). To determine whether this upregulation reflects a cell-intrinsic effect, we co-adoptively transferred congenically marked naïve WT and *Etv4* KO OT-I cells into B16-OVA tumors. Under these conditions, T-bet expression remained elevated in *Etv4* KO CD8^+^ T cells (Fig. 6c), confirming that the effect is cell-intrinsic. Next, we investigated whether ETV4 directly regulates *Tbx21* transcription. In CD8^+^ T cells subjected to chronic TCR stimulation, we found that ETV4 expression peaked at the second activation cycle and declined during subsequent stimulations (Fig. 6d), reflecting its dynamic regulation during T cell exhaustion. ChIP–qPCR performed after the second round of stimulation revealed direct binding of ETV4 to *Tbx2*1 promoter regions containing ETV4-binding motifs (Fig. 6e, f). Together, these data suggest that loss of ETV4 results in derepression of T-bet expression, thereby contributing to elevated IFNγ production in *Etv4* KO CD8^+^ T cells.

**Figure 6.**
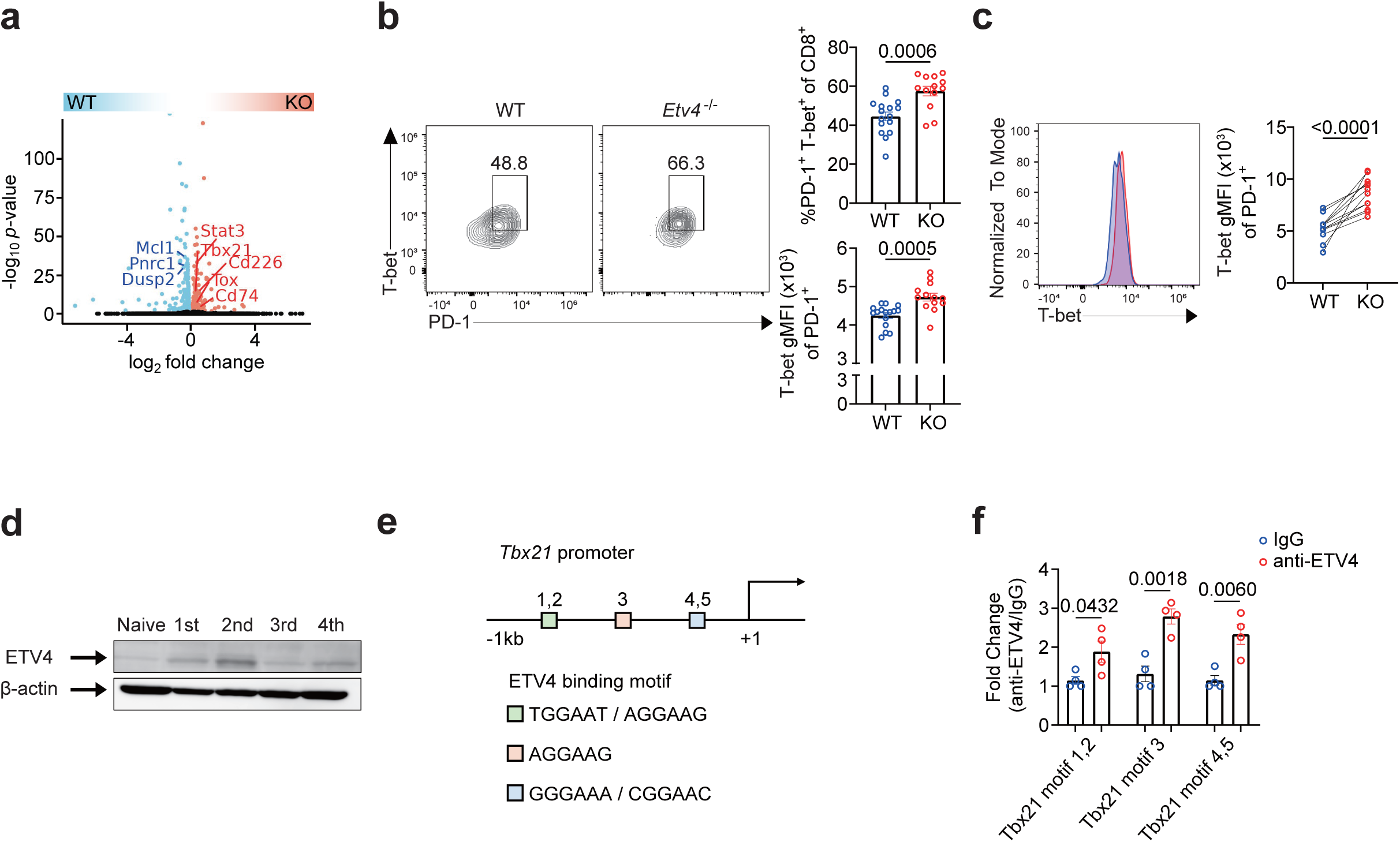
ETV4 suppresses T-bet expression in Tex cells. **a,** Volcano plot of differentially expressed genes between WT and *Etv4* KO CD8^+^ T cells. **b,** Representative flow cytometry plots and percentage of PD-1^+^T-bet^+^ CD8^+^ T cells and gMFI quantification of T-bet expression in PD-1^+^ CD8^+^ T cells infiltrating MC38 tumors in WT (n=16) and *Etv4* KO (n=13) mice. **c,** Representative histograms and gMFI quantification of T-bet expression in PD-1^+^ OT-I cells from B16-OVA tumor-bearing *Rag1* KO mice after intratumoral co-adoptive transfer of CD45.1 WT (blue) and CD45.2 *Etv4* KO (red) (n=12) OT-I cells. **d,** Western blot analysis of ETV4 expression profiles in CD8^+^ T cells during *in vitro* T cell exhaustion models. **e,** Schematic of the *Tbx21* promoter region with three predicted ETV4-binding motifs**. f,** ChIP-qPCR of *Tbx21* promoter regions containing ETV4-binding motifs in CD8^+^ T cells after two rounds of continuous TCR stimulation, using IgG or anti-ETV4 antibody (n=4). Data represent three independent experiments for **b-d** and **f**. Error bars indicate mean ± s.e.m. Statistical significance was determined using unpaired two-tailed Student’s t-tests for **b** and **f**, and paired two-tailed Student’s t-tests for **c**. Data in **b** and **f** were pooled from two independent experiments.

### CD74 is a conserved functional marker across human cancers

Finally, we investigated whether the role of CD74 that we had observed in mice is conserved in humans. To this end, we re-analyzed public human pan-cancer datasets encompassing eight tumor types^50^ (Fig. 7a, b and Extended Data Fig. 8a, b). The analysis was restricted to CD8^+^ T cells isolated from blood, normal tissue, and tumor samples, allowing a comprehensive comparison of expression patterns across tissue contexts. In human CD8^+^ T cells, *CD74* expression positively correlated with exhaustion-associated genes, such as *HAVCR2*, *LAG3*, *CTLA4*, *TIGIT*, *TOX*, *ENTPD1*, and *CXCL13*, but not with Tpex-related genes, including *CCR7*, *IL7R*, and *TCF7* (Fig. 7c). In line with these findings, analysis of murine tumor-infiltrating CD8^+^ T cells showed that CD74 was preferentially expressed in TIM3⁺ and TIM3^hi^ subsets (Extended Data Fig. 9a, b), although its geometric mean fluorescence intensity (gMFI) was comparable between Tpex and Tex cells (Fig. 3d, e).

**Figure 7.**
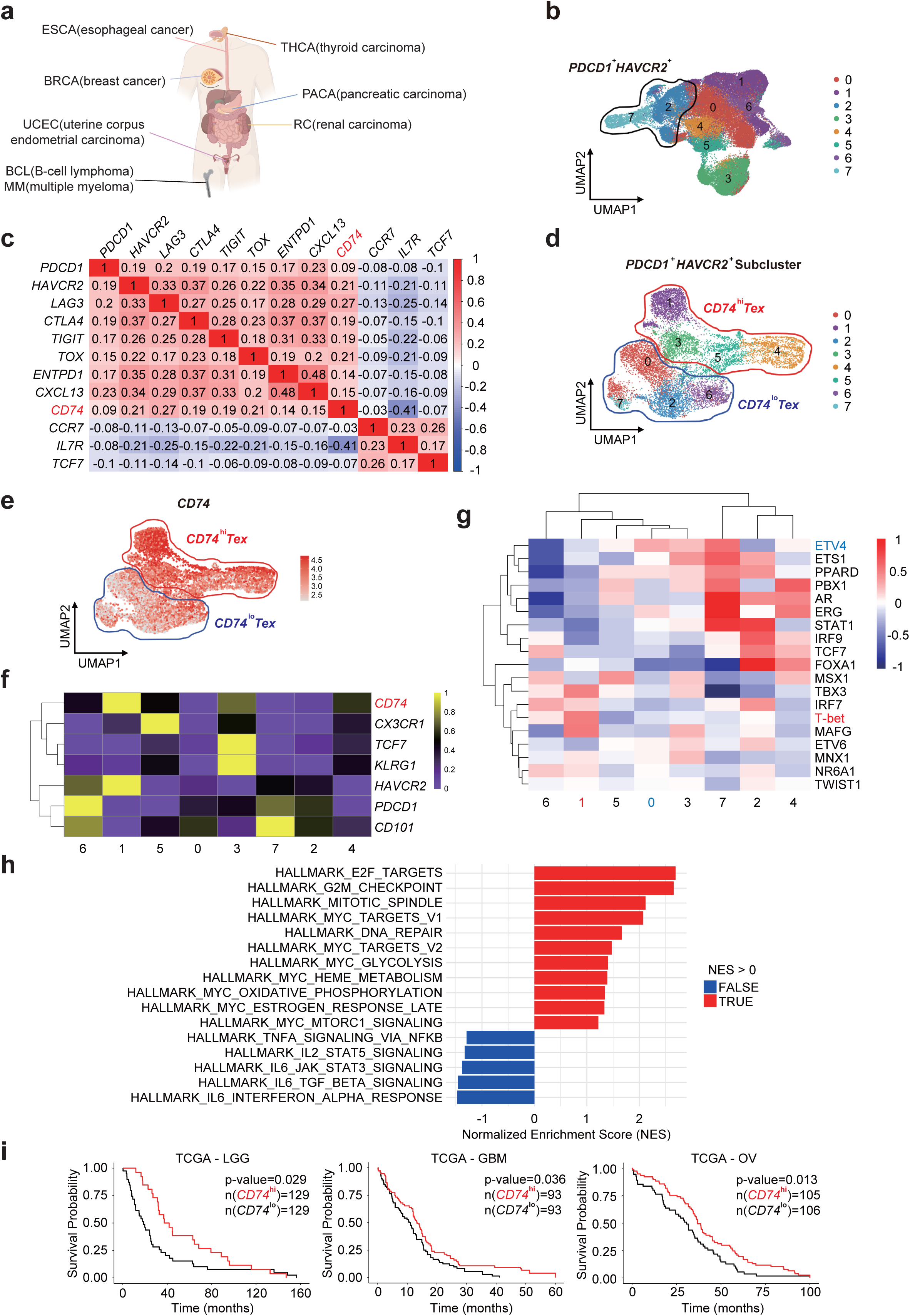
CD74 is a conserved functional marker across human pan-cancer. **a,** Schematic of eight human tumor types included in the CD8^+^ T cell scRNA-seq pan-cancer analysis^50^. **b,** UMAP dimensional reduction of scRNA-seq data from blood, normal tissue, and tumor-infiltrating CD8^+^ T cells. **c,** Heatmap depicting correlations between genes associated with exhaustion and Tpex signatures. **d–h,** UMAP visualization and functional characterization of Tex (*PDCD1*⁺ *HAVCR2*⁺) cell subclusters: UMAP visualization of subclusters **(d)**, feature plot of *CD74* expression (red) **(e)**, heatmap of exhaustion marker expression **(f)**, heatmap of various transcription factor activities **(g)**, and GSEA bar plot comparing *CD74*^hi^ versus *CD74*^lo^ (*PDCD1*⁺*HAVCR2*⁺) Tex cells, with red and blue bars denoting upregulated and downregulated pathways, respectively **(h)**. **i,** TCGA survival analysis comparing patients with *CD74*^hi^ and those with *CD74*^lo^ normalized expression in lower-grade glioma (LGG), glioblastoma (GBM), and ovarian cancer (OV). *CD74* expression in each tumor sample was normalized to *CD8A* expression to control for variability in CD8^+^ T cell infiltration.

To further assess functional features of CD74 and its relationship with ETV4, we subclustered *PDCD1*^+^*HAVCR*2^+^ Tex cells (Fig. 7d and Extended Data Fig. 8a, b). Notably, cluster 1, which had the highest *CD74* expression levels, exhibited low ETV4 but high T-bet activity, whereas cluster 0, which had the lowest *CD74* expression levels, showed the opposite pattern (Fig. 7e-g). This suggests that *CD74* expression in human Tex cells is positively correlated with T-bet activity and negatively correlated with ETV4 activity. Consistent with increased T-bet activity^51, 52, 53, 54^, GSEA revealed that *CD74*^hi^ Tex cells were enriched for pathways related to proliferation and metabolism (Fig. 7h). Conversely, pathways downregulated in *CD74*^hi^ Tex cells included multiple cytokine signaling cascades, such as TNF^55^, IL-2^56^, IL-6–STAT3^57^, TGF-β^58^, and IFNα signaling^59,60^, all of which are known to impair CD8^+^ T cell function. The negative enrichment of these exhaustion-inducing pathways supports a functionally active phenotype for *CD74*^hi^ Tex cells.

Finally, we examined the association between *CD74* expression and clinical outcomes in cancers with poor prognosis. Re-analysis of The Cancer Genome Atlas (TCGA) RNA-seq data from lower-grade glioma (LGG), glioblastoma (GBM), and ovarian cancer—tumors in which immunotherapies such as anti–PD-1 have shown limited efficacy owing to immunosuppressive extrinsic factors against CD8^+^ T cell immunity^61, 62^—revealed that higher normalized *CD74* expression levels correlated with improved survival (Fig. 7i). Collectively, these observations support CD74 as a candidate marker of functional Tex cells across human cancers, with potential relevance for patient prognosis.

## Discussion

T cell exhaustion remains a major challenge in cancer immunotherapy. However, this state is not uniformly detrimental. Tpex cells lack immediate cytotoxic capacity, whereas Tex cells lose their proliferative potential and fail to sustain long-term immunity^4^. Thus, maintaining a balanced differentiation between these two subsets is essential for durable and effective anti-tumor responses. Certain strategies involve targeting the transcription factor TOX; however, these result in the depletion of both Tpex and Tex cells^10, 40^.

In this context, our study has identified ETV4 as a promising target to enhance anti-tumor immunity without disrupting the balance between Tpex and Tex cells. *Etv4* deletion does not affect the Tpex pool (Fig. 2c, f), whereas it promotes Tex differentiation and enhances cytotoxicity, particularly through the expansion of CD74^+^ Tex cells with superior effector function. Although CD74 interacts with MIF, typically regarded as immunosuppressive^63^, its restricted expression in Tex-KLR cells and MIF-independent cell-intrinsic function support its contribution to enhanced T cell functionality. Therefore, targeting ETV4 in chimeric antigen receptor-T cells to enhance CD74 expression may be a potential strategy to improve therapeutic efficacy. In contrast, direct targeting of CD74 with agonistic antibodies may entail risks. CD74 mediates immunosuppressive signaling via MIF in macrophages and DCs, and systemic activation of this pathway could undermine T cell responses, particularly given the abundance of myeloid cells in the TME^39^. Thus, therapeutic strategies involving CD74 should aim to selectively modulate CD8^+^ T cells while minimizing effects on myeloid populations.

Another important implication of our findings is the potential use of CD74 as a functional marker. Commonly used functional Tex markers such as CX3CR1 are often underrepresented in scRNA-seq datasets (Extended Data Fig. 8a, b), limiting their utility to be used to reliably distinguish Tex subsets. In contrast, CD74 is consistently and robustly detected across various scRNA-seq platforms (Extended Data Fig. 8a, b), enabling more precise classification of Tex populations. In line with our findings, other scRNA-seq datasets also report high *Cd74* expression in Tpex cells^64^. Interestingly, in our *in vitro* chronic TCR stimulation model, CD74 expression was higher in Tpex cells than in Tex cells (Extended Data Fig. 9c), suggesting that CD74 expression may vary depending on microenvironmental contexts. As Tpex populations segregate according to CD74 expression (Fig. 3d, e and Extended Data Fig. 9d, e), we speculate that this distinction reflects two lineages with divergent developmental trajectories: CD74^+^ Tpex cells may give rise to functionally superior CD74^+^ Tex cells, whereas CD74^−^ Tpex cells may differentiate into dysfunctional CD74^−^ Tex cells. This model raises the possibility that functional heterogeneity within Tex cells originates as early as the progenitor stage. Further investigations will be required to delineate how CD74 regulates Tpex fate decisions and contributes to Tex diversity.

The molecular mechanism by which CD74 influences the fate of Tex cells remains unclear. Although the intracellular domain of CD74 can act as a transcriptional regulator, its formation requires ligand–receptor interactions^63, 65^. Data from our *in vitro* studies suggest that MIF does not signal through CD74 in CD8^+^ T cells, implying the involvement of alternative ligands. Our TCGA survival analyses revealed that higher *CD74* expression correlates with improved patient survival in GBM and LGG (Fig. 7i). In GBM, amyloid precursor protein (APP) has been identified as a CD74 ligand on tumor-associated macrophages (TAMs), where the APP–CD74 axis suppresses phagocytosis and promotes tumor progression^66^. We speculate that elevated CD74 expression on CD8^+^ T cells may competitively bind APP, thereby mitigating its inhibitory effects on TAM phagocytosis and indirectly enhancing anti-tumor immunity.

In summary, our study identifies ETV4 as a promising therapeutic target. This strategy will potentially enhance the effector function of Tex cells and promote the differentiation of cytotoxic CD74^+^ Tex cells, thereby circumventing the limitations of TOX inhibition. Our findings challenge the view of exhaustion as a fixed or irreversible state and instead suggest that it represents a modifiable differentiation program. Although an overabundance of Tpex cells can limit cytolytic activity and expanded Tex cells often exhibit dysfunction, restoring Tex functionality by modulating ETV4 or CD74 may elicit robust tumoricidal responses and ultimately improve clinical outcomes.

## Materials and Methods

### Mice

All mice were maintained on a C57BL/6 background. Whole-body *Etv4* KO mice were generated using the CRISPR/Cas9 system (Korea Mouse Phenotyping Center, Seoul, Korea). Tg(TcraTcrb)1100Mjb/J (OT-I), *Rag1* KO, and CD45.1 mice were obtained from the Jackson Laboratory. All experiments were conducted with 6–9-week-old mice unless otherwise indicated. Animals were maintained in a specific pathogen-free animal facility under a standard 12-h light/12-h dark cycle. Mice were fed standard rodent chow and water *ad libitum*. Each experiment was performed using 5–15 mice that were assigned randomly to the experimental groups. Male and female mice were also randomly assigned to the experimental groups. All procedures were approved by the Pohang University of Science and Technology Institutional Animal Care and Use Committee.

### Cell lines and tumor growth curves

MC38 (CVCL_B288) and B16-OVA (CVCL_WM77) cells were obtained from Sigma-Aldrich. The MC38 and B16-OVA cell lines were confirmed to be mycoplasma-free. All immortalized cell lines were maintained in laboratory-made DMEM medium (DMEM with 10% fetal bovine serum and 5% penicillin plus streptomycin). The cultures were incubated in temperature-stable and partial pressure–stable conditions at 37 °C and 5% CO_2_. WT, *Etv4* KO, and *Rag1* KO mice were injected with 2×10⁵ MC38 cells (subcutaneously) or B16-OVA cells (intradermally) suspended in pure DMEM into the right flank. The tumors were measured every 3 days using digital calipers. The maximal size to which tumors were allowed to grow in this study was 15 mm in any direction. Whenever the maximal tumor size was reached, the tumor-bearing mouse was sacrificed. Mice were excluded from analysis if tumors became ulcerated or failed to reach the maximal size.

### Tumor collection and mechanical disruption

For tumor-infiltrating lymphocyte (TIL) analysis, tumors were harvested on days 12–14. For immune cell isolation, whole tumors were dissected using a razor blade and incubated in digestion buffer (PBS containing collagenase D [0.90 mg/mL] and DNase I [0.19 mg/mL]) at 37 °C for 30 min. Following enzymatic digestion, tumors were mechanically dissociated using the back end of a 5-mL syringe plunger and filtered through 70-µm cell strainers (cat# 93070, SPL). Immune cells were then enriched using OptiPrep™ Density Gradient Medium (cat# D1556-250ML, Sigma-Aldrich) according to the manufacturer’s instructions.

### CD8^+^ T cell negative selection

For *in vitro* and adoptive transfer experiments, naïve CD8⁺ T cells were purified to >95% purity from pooled lymph nodes and spleens of WT, *Etv4* KO, OT-I, and *Etv4* KO OT-I mice. Cells were isolated using a modified CD4⁺ negative selection protocol (Stemcell Technologies) with The Big Easy™ EasySep™ Magnet, EasySep™ Mouse Streptavidin RapidSpheres™ Isolation Kit (cat# 19860, Stemcell Technologies), and a custom mixture of biotinylated antibodies (BioLegend). CD4 (RM4-5, cat# 100508), CD11b (M1/70, cat# 101204), CD11c (N418, cat# 117304), CD19 (6D5, cat# 115504), and B220 (RA3-6B2, cat# 103204) were each added at 4 µl per sample unit. CD49b (DX5, cat# 108904), TER119 (TER-119, cat# 116204), Gr-1 (RB6-8C5, cat# 108404), and TCR γδ (UC7-13D5, cat# 13-5811-85) were each added at 3 µL per sample unit. The total antibody mixture (excluding CD44) was adjusted to 50 µL per sample. All steps were performed at room temperature. EasySep™ Mouse FcR Blocker (cat# 18731, Stemcell Technologies) was added at 50 µL per sample and incubated for 10 min. Subsequently, the biotinylated antibody cocktail (excluding CD44) was added and incubated for 12 min. CD44 (IM7, cat# 103003) was subsequently added at 1.3 µL per sample and incubated for 1 min 40 sec. EasySep™ Streptavidin RapidSpheres™ (cat# 50001, Stemcell Technologies) were then added at 40 µL per sample and incubated for 2 min 30 sec. After incubation, 5 mL of FACS buffer (2% FBS in PBS) was added, and the tubes were placed on The Big Easy™ Magnet for 2 min 30 sec. The untouched naïve CD8⁺ T cell fraction was collected and used for downstream applications.

### *In vitro* CD8^+^ T cell cultures

To mimic CD8⁺ T cell exhaustion, we adapted previously published protocols^12^. Naïve CD8⁺ T cells were isolated from pooled lymph nodes and spleens of mice using a negative selection method. For each T cell activation, Mouse CD3ε NALE (145-2C11, cat# 553057, BD Pharmingen) and Mouse CD28 NALE (37.51, cat# 553294, BD Pharmingen) were pre-coated onto immunoplates (cat# 32296, SPL) at a 2:1 ratio and incubated overnight at 4 °C. Cells were first activated for 3 days in the presence of IL-2 (cat# 212-12, Peprotech, 10 ng/mL) and IL-12 (cat# 419-ML-010/CF, R&D Systems, 10 ng/mL). This was followed by a second activation for 1 day with IL-2 (10 ng/mL) alone. The third and fourth rounds of activation were performed for 2 days each with IL-2 (10 ng/mL). For co-culture experiments, CD8⁺ T cells from CD45.1, CD45.1/2, or CD45.2 WT mice (as indicated in each experiment) and CD45.2 *Etv4* KO mice were mixed at a 1:1 ratio, with the actual input ratio carefully controlled such that neither population exceeded approximately 58% of the total (i.e., within 42:58). A total of 1×10⁵ cells were seeded per well. For IFNγ (cat# 575304, BioLegend, 10 ng/mL) treatment, the cytokine was included throughout all four rounds of activation. For experiments with anti-IFNγ (cat# 505802, BioLegend, 5 µg/mL), the antibody was added during the third activation cycle only. For MIF (cat# 1978-MF-025/CF, R&D systems, 100 ng/mL) treatment, the cytokine was included after transduction.

### Flow sorting and cytometry

Flow sorting and cytometry studies were performed as previously described^67^, with minor modifications. Viability staining with Ghost Dye™ (cat# 13-0870, Tonbo, 1:1000 dilution) and surface staining (1:200 dilution) were performed in FACS buffer at 4 °C for 30 min. Following viability and surface staining, cells were subject to intracellular staining for transcription factors or cytokines. For intracellular staining of transcription factors (1:100 dilution), cells were permeabilized using the Foxp3/Transcription Factor Staining Kit (cat# 00-5523-00, eBioscience) according to the manufacturer’s protocol, washed in 1× permeabilization washing buffer, and stained in the same buffer at 4 °C for 40 min. For intracellular cytokine staining (1:100 dilution), cells were fixed and permeabilized using the BD Cytofix/Cytoperm Fixation/Permeabilization Kit (cat# 554714, BD Biosciences) according to the manufacturer’s protocol, washed in 1× permeabilization washing buffer, and stained in permeabilization buffer at 4 °C for 40 min. For intracellular staining of transduced GFP⁺ cells, cells were fixed in 4.2% paraformaldehyde at 4 °C for 30 min, followed by intracellular staining of transcription factors or cytokines. The following antibodies or staining reagents were purchased from BioLegend: CD45 (30-F11, cat# 103130), CD45.1 (A20, cat# 110724), CD45.2 (104, cat# 109846), CD8α (53-6.7, cat# 100714), CD4 (RM4-5, cat# 100552), CD279/PD-1 (29F.1A12, cat# 135220), CD366/Tim-3 (RMT3-23, cat# 119723), CD223/LAG-3 (C9B7W, cat# 125210), CD39 (Duha59, cat# 143803), CX3CR1 (SA011F11, cat# 149016), KLRG1/MAFA (2F1/KLRG1, cat# 138414), CD74 (In1/CD74, cat# 151004), Annexin V (cat# 640919), CD152/CTLA-4 (UC10-4B9, cat# 106305), T-bet (4B10, cat# 644832), TNF (MP6-XT22, cat# 506341), IFNγ (XMG1.2, cat# 505825), Granzyme B (QA18A28, cat# 396406), CD11b (M1/70, cat# 101223), Ly6C (HK1.4, cat# 128007), NK1.1 (PK136, cat# 108710), and F4/80 (BM8, cat# 123145). TOX (TXRX10, cat# 12-6502-82), Eomes (Dan11mag, cat# 12-4875-80), Ki-67 (SolA15, cat# 11-5698-80), CD11c (N418, cat# 47-0114-80), and Ly6G (RB6-8C5, cat# 17-5931-81). CD19 (1D3, cat# 562701), CD25 (PC61, cat# 561112), Stat3 (pY705) (4/P-STAT3, cat# 557815), and CD44 (IM7, cat# 561860) were purchased from BD Biosciences. FOXP3 (FJK-16s, cat# 48-5773-82) was obtained from Invitrogen. TCF-1 (C63D9, cat# 14456S) was obtained from Cell Signaling Technology. Flow cytometry was performed using a Beckman Coulter CytoFLEX LX (Ⅱ). Data were analyzed via FlowjoV10.

### CD8^+^ T cell cytokine production assay

*In vitro*-activated CD8⁺ T cells or CD8⁺ T cells from MC38 tumors were stimulated with phorbol 12-myristate 13-acetate and ionomycin for 3 h 30 min in RPMI medium supplemented with BD GolgiPlug™ Protein Transport Inhibitor (containing Brefeldin A; cat# 555029, BD Biosciences; 1:1000 dilution) and BD GolgiStop™ Protein Transport Inhibitor (containing Monensin; cat# 554724, BD Biosciences; 1:1500 dilution). OT-I cells from B16-OVA tumors were stimulated with OVA 257-264 peptide (H-2Kᵇ-restricted OVA MHC class I epitope; cat# vac-sin, InvivoGen; 1 µg/mL) for 6 h in RPMI medium supplemented with the same concentrations of GolgiPlug™ and GolgiStop™.

### Plasmids production

The coding sequence of mouse *Cd74* was amplified from cDNA of MC38 using Pfu-X DNA polymerase (cat# SPX16, SolGent) and cloned into the MIGR1-GFP vector (cat# 27490, Addgene). The C-terminal region of CD74 fused with a FLAG tag was cloned into the MIGR1-GFP vector. The primers used for cloning was as follows: CD74 Forward; 5′-CCCTCGAGATGGATGACCAACGCGACCTCATCT-3′, CD74 Reverse; 5′-CGGAATTCCTTGTCATCGTCGTCCTTGTAGTCCAGGGTGACTTGACCCA-3′.

The shRNA expression vector for the knockdown of mouse *Cd74* was generated using the MSCV-LTRmiR30-PIG (LMP) vector (Open Biosystems) in accordance with the manufacturer’s instructions. The target sequences were as follows: for sh*Cd74*-1; 5′-CCCAGAACCTGCAACTGGA-3′, for sh*Cd74*-2; 5′-CGTCCAATGTCCATGGATA-3′, for sh*Cd74*-3; 5′-CAGAGAATCTGAAGCATCT-3′.

### Retroviral transduction

The vector was transfected into Plat-E retroviral packaging cells. Retroviral supernatants were collected 48 and 72 h post-transfection, concentrated using the Retro-X concentrator (cat# 631456, Takara) according to the manufacturer’s instructions, and stored in RPMI medium. OT-I cells were isolated using the negative selection method described above and stimulated for 24 h with 2 μg/mL αCD3ε and 1 μg/mL αCD28. Cells were then pelleted and transduced with the viral supernatant in the presence of 5μg/mL polybrene by centrifugation at 2,000 rpm for 2 h. Transduced cells were cultured for 4 days for expansion with IL-2 (cat# 212-12, Peprotech, 10 ng/mL) and subsequently sorted based on GFP expression. After sorting, GFP⁺ cells were further expanded for further experiments.

### Intratumoral adoptive transfer

For tumor growth analysis, *Rag1* KO, CD45.1, and CD45.2 mice were intradermally inoculated with B16-OVA tumor cells (2×10⁵ cells). When tumor volumes reached 80–250 mm³, *Rag1* KO mice received intratumoral injections of pre-activated (2-day) WT CD45.1 OT-I cells or *Etv4* KO CD45.2 OT-I cells (2×10⁶ cells each). For TIL analysis comparing WT CD45.1 and *Etv4* KO CD45.2 OT-I cells, a similar protocol was used, except that the mice received a 1:1 mixture of naïve WT CD45.1 and naïve *Etv4* KO CD45.2 OT-I cells (1×10⁶ total; 0.5×10⁶ WT + 0.5×10⁶ *Etv4* KO per mouse) to track their bona fide activation within the TME. For tumor growth analysis involving EV and *Cd74* KD OT-I cells, CD45.1 immunocompetent mice bearing B16-OVA tumors (80–250 mm³) received intratumoral injections of pre-activated (1-day) EV CD45.2 OT-I cells or *Cd74* KD CD45.1/2 OT-I cells (2×10^6^ cells each). For TIL analysis in this context, mice were injected with a 1:1 mixture of pre-activated (1-day) EV CD45.2 and *Cd74* KD CD45.1/2 OT-I cells (1×10⁶ total; 0.5×10⁶ EV + 0.5×10⁶ *Cd74* KD per mouse).

### Western blotting

Chronically activated CD8^+^ T cells were lysed in RIPA buffer (50 mM Tris-HCL pH 7.4, 1 mM PMSF, 1x Roche Complete Protease Inhibitor Cocktail, 0.5% sodium deoxycholate, 0.1% SDS, 150 mM NaCl). Protein concentrations were measured using a BCA kit (cat# 23225, Pierce). Equal amounts of protein samples were separated using 10% SDS-PAGE and transferred onto nitrocellulose membranes (cat# 1620115, Bio-Rad). The following primary antibodies were used: anti-ETV4 (cat# 10684-1-AP, Proteintech) and anti-β actin (cat# sc-47778, Santa Cruz Biotechnology). Membranes were incubated with secondary antibodies conjugated to horseradish peroxidase and developed using Clarity Western ECL substrates (cat# 1705061, Bio-Rad). Images were acquired using an ImageQuant LAS 500 instrument (GE Healthcare).

### RT-qPCR

RT-qPCR was performed as previously described^68^, with minor modifications. Total RNA was purified using the RiboEx reagent, and 1.5 μg of total RNA was reverse-transcribed into cDNA using a GoScript Reverse Transcription System (cat# A5004, Promega) as per manufacturer’s instructions. The SYBR Green real-time PCR master mix (cat# TOQPK-201, Toyobo) was used for qPCR. Gene expression was normalized to 18S rRNA levels. Primer sequences are listed in Supplementary Table S1.

### ChIP-qPCR

Naïve CD8⁺ T cells (1×10⁷) underwent two consecutive rounds of continuous activation and were subsequently cross-linked with 1% paraformaldehyde for 15 min under constant agitation. The reaction was quenched with 1 M glycine for 5 min, and cells were washed twice with cold PBS and centrifuged at 4200 rpm for 5 min. The pellet was resuspended in Buffer 1 (50 mM HEPES-KOH, pH 7.5; 140 mM NaCl; 1 mM EDTA, pH 8.0; 10% glycerol; and 0.5% NP-40) and rotated at 4 °C for 10 min to lyse nuclei. After a 5-min centrifugation at 4200 rpm, the lysate was resuspended in Buffer 2 (10 mM Tris-HCl, pH 8.0; 300 mM NaCl; 0.1% sodium deoxycholate; 1% Triton X-100; 1 mM EDTA, pH 8.0; and 0.5 mM EGTA, pH 8.0) and sonicated (90 cycles of 20 sec on/40 sec off) to shear DNA. The supernatant was collected after centrifugation at 9400 *g* for 10 min at 4 °C, and 5% was stored at −80 °C as input. For immunoprecipitation, chromatin was precleared with protein G agarose (cat# 16-266, Merck) for 90 min in Buffer 2, followed by incubation overnight at 4 °C with either 2 µg of normal rabbit IgG (cat# 2729, Cell Signaling Technology) or anti-ETV4 antibody (cat# 10684-1-AP, Proteintech). The chromatin–antibody complexes were captured using protein G agarose for an additional 4 h at 4 °C. Subsequently, the samples were washed sequentially with low-salt buffer (4 mM EDTA, 2% Triton X-100, 40 mM Tris-HCl, pH 8.0, 300 mM NaCl, 0.2% SDS), high-salt buffer (4 mM EDTA, 2% Triton X-100, 40 mM Tris-HCl, pH 8.0, 1 M NaCl, 0.2% SDS), LiCl buffer (0.5 M LiCl, 20 mM Tris-HCl, pH 8.0, 2 mM EDTA, 2% NP-40, 2% sodium deoxycholate), and TE buffer (20 mM Tris-HCl, pH 8.0, 2 mM EDTA). Bound chromatin was eluted using elution buffer (0.1 M NaHCO₃ and 0.5% SDS) and reverse-crosslinked with 200 mM NaCl at 65 °C overnight. RNA and proteins were subsequently digested with RNase A and protease K, respectively, and DNA was purified using the Expin CleanUp SV kit (cat# 113-150, GeneAll). Finally, qPCR was performed to quantify the relative enrichment of *Tbx21* promoter regions in the immunoprecipitated DNA fragments. qPCR primer sequences used are listed in Supplementary Table S1.

### Analysis of public single-cell RNA-seq datasets

Public scRNA-seq datasets from murine blood and MC38 tumor-infiltrating CD8⁺ T cells, as well as from human blood, normal tissues, and tumors, were used to evaluate the expression of *Etv4*/*ETV4* and *Cd74*/*CD74* across CD8⁺ T cell subsets. The MouseIntegratedBlood Seurat object was obtained from the Gene Expression Omnibus (GSE158520)^31^, and the human pan-cancer count matrix was obtained from the Gene Expression Omnibus (GSE156728)^50^. CD8⁺ T cells were computationally isolated from other immune populations, and the top 2,000 highly variable genes (HVGs) were selected for dimensionality reduction via principal component analysis (PCA). The optimal number of principal components was determined by the inflection point on the elbow plot and subsequently used for uniform manifold approximation and projection (UMAP) and Louvain-based clustering. The expression and transcription factor activity of *Etv4*/*ETV4*, as well as *Cd74*/*CD74* expression, were then assessed across blood, normal tissue, and tumor tissue clusters and visualized using heat maps. Transcription factor activity for each cluster was inferred using the run_ulm function of the decoupleR (v2.9.7) R package^69^. For gene set enrichment analysis (GSEA) in human pan-cancer scRNA-seq datasets, differentially expressed genes (DEGs) between *CD74*^hi^ Tex and *CD74*^lo^ Tex were ranked by average log_2_ fold change and subjected to GSEA with the Molecular Signatures Database (MSigDB) using fgsea (v1.28.0)^70^ and msigdbr (v7.5.1)^71^ R packages.

### Single-cell RNA-sequencing

Single-cell RNA sequencing libraries were generated using the Chromium Next GEM Single Cell 3’ Kit v3.1 (10x Genomics, PN-1000269) together with the Dual Index Kit TT Set A (PN-1000215). Cell suspensions were loaded onto a Chromium Next GEM Chip G (PN-2000127) and processed on a Chromium iX instrument to generate gel bead-in-emulsions (GEMs), targeting a recovery of 10,000 cells per channel. GEMs were subjected to reverse transcription in a thermal cycler, and the resulting cDNA was purified and amplified. Library preparation was performed according to the manufacturer’s protocol, including enzymatic fragmentation, end repair, A-tailing, adaptor ligation, and sample index PCR. cDNA and library quality were evaluated using a High Sensitivity DNA ScreenTape Analysis kit on a TapeStation system (Agilent). Libraries were pooled and sequenced on an Illumina NovaSeq 6000 platform with 100-bp paired-end reads, aiming for a minimum depth of 20,000 read pairs per cell for 3′ gene expression libraries.

### WT vs *Etv4* KO CD8^+^ T cells scRNA-seq data preprocessing

Raw reads for scRNA-seq were aligned to the mouse reference genome (GRCm39) using CellRanger (v8.0.1) ^72^ with the Ensembl GRCm39.110 GTF annotation. Empty droplets were identified and removed using the emptyDrops function of the DropletUtils (v1.22.0) R package^73^. Low-quality cells using detected unique molecular identifiers (UMIs) lower than 700 or with a percentage of genes mapped to mitochondrial genes higher than 10% were filtered out using the addPerCellQC function in scater (v1.30.1) R package^74^. To normalize cell-specific biases, cells were grouped using the quickCluster function in scran (v1.30.0) R package^75^ and cell-specific size factors were calculated using the computeSumFactors function of the same package. Raw unique molecular identifier counts were normalized by cell-specific size factor and log_2_-transformed with the pseudocount of 1 using the logNormCounts function of scater. HVGs were identified using the modelGeneVar function of scran with false discovery rate (FDR) < 0.05. For downstream analysis, the normalized counts matrix was scaled and PCA was performed with HVGs. All cells were clustered using the FindClusters function of the Seurat (v5.0.3) R package^76^ on the top 18 principal components (PCs) with a resolution of 0.8. Cells were visualized on a two-dimensional UMAP plot using the RunUMAP function of Seurat. The scDblFinder (v1.16.0) R package^77^ was applied to detect putative doublets. These doublets were manually inspected to determine whether they expressed two or more lineage-specific markers and were removed if confirmed. Cells annotated as myeloid cells, stromal cells and CD4^+^ T cells were removed for further analysis. The remaining cells were re-processed using the same methods described above. CD8^+^ T cells and Tex cells were clustered and visualized using the top 12 and 10 PCs, respectively.

### WT vs *Etv4* KO CD8^+^ T cells scRNA-seq data analysis

For the PD-1^+^ CD8^+^ T cell subtype, composition differences between WT and *Etv4* KO were assessed using odds ratios and 95% confidence intervals. *P* values, calculated by Fisher’s exact test, were adjusted for multiple testing by the Benjamini-Hochberg method. Gene set signature scores for each cell were calculated using the AddModuleScore function of Seurat. DEGs between WT and *Etv4* KO Tex cells were identified using the FindMarkers function of Seurat. The DEG list is provided in Supplementary Table S2. Transcription factor activity for each cluster was inferred using the run_ulm function of the decoupleR (v2.9.7) R package^69^. For GSEA, DEGs between WT and *Etv4* KO in cluster 1 of Tex cells were identified using the FindMarkers function. The DEG list of cluster 1 is provided in Supplementary Table S3. The DEGs were ranked by average log_2_ fold change and subjected to GSEA with the MSigDB using fgsea (v1.28.0)^70^ and msigdbr (v7.5.1)^71^ R packages.

### TCGA data analysis

RNA-seq and clinical data of TCGA glioblastoma, lower-grade glioma, and ovarian cancer cohorts were obtained from the National Cancer Institute Genomic Data Commons Data Portal using the R package *TCGAbiolinks* (version 3.20.0). Raw read counts (“HTSeq-Counts”) were transformed into transcripts per million, and *CD74* expression values were normalized to *CD8A* to account for inter-patient variability in CD8⁺ T cell infiltration within tumors. Patients were stratified into *CD74*^hi^ (top 25%) and *CD74*^lo^ (bottom 25%) groups based on normalized *CD74* expression. Survival analyses were performed using the Kaplan–Meier method, and statistical significance was evaluated with the log-rank test.

### Statistical analysis

All experiments were performed at least in duplicate, independently. Datasets were analyzed using the two-tailed unpaired, paired Student’s *t*-test and two-way ANOVA with Tukey’s multiple comparison using the Prism 9.0 software (GraphPad). A P-value < 0.05 was considered significant. In the bar graphs, bars and error bars indicate mean ± standard error of the mean.

## Data availability

The raw WT vs *Etv4* KO CD8^+^ T cells scRNA-seq data generated in this study have been deposited in the NCBI Sequence Read Archive (SRA) database under accession code PRJNA1328826. All other data are available in the main text or supplementary materials.

## Acknowledgments

We thank the members of Lee laboratory for their inputs and comments on the study. This study was supported by the National Research Foundation (NRF) of Korea grants funded by the Korean government (RS-2021-NF000572, RS-2023-00260454, RS-2024-00336114, RS-2025-02216523, and RS-2025-19032970) and a Korea Basic Science Institute (National Research Facilities and Equipment Center) grant funded by the Ministry of Education (2021R1A6C101A390). MK, YH, YS, and JSP received support from the BK21 Program.

## Author contributions

Conceptualization: DWL; Methodology: DWL; Investigation: DWL, YWK, HMG, YHC, and JYL; Writing—original draft: DWL; Writing—review & editing: DWL and YL; Visualization: DWL and HMG; Supervision: YL; Funding acquisition: YL

---

**Extended Data Figure 1.**
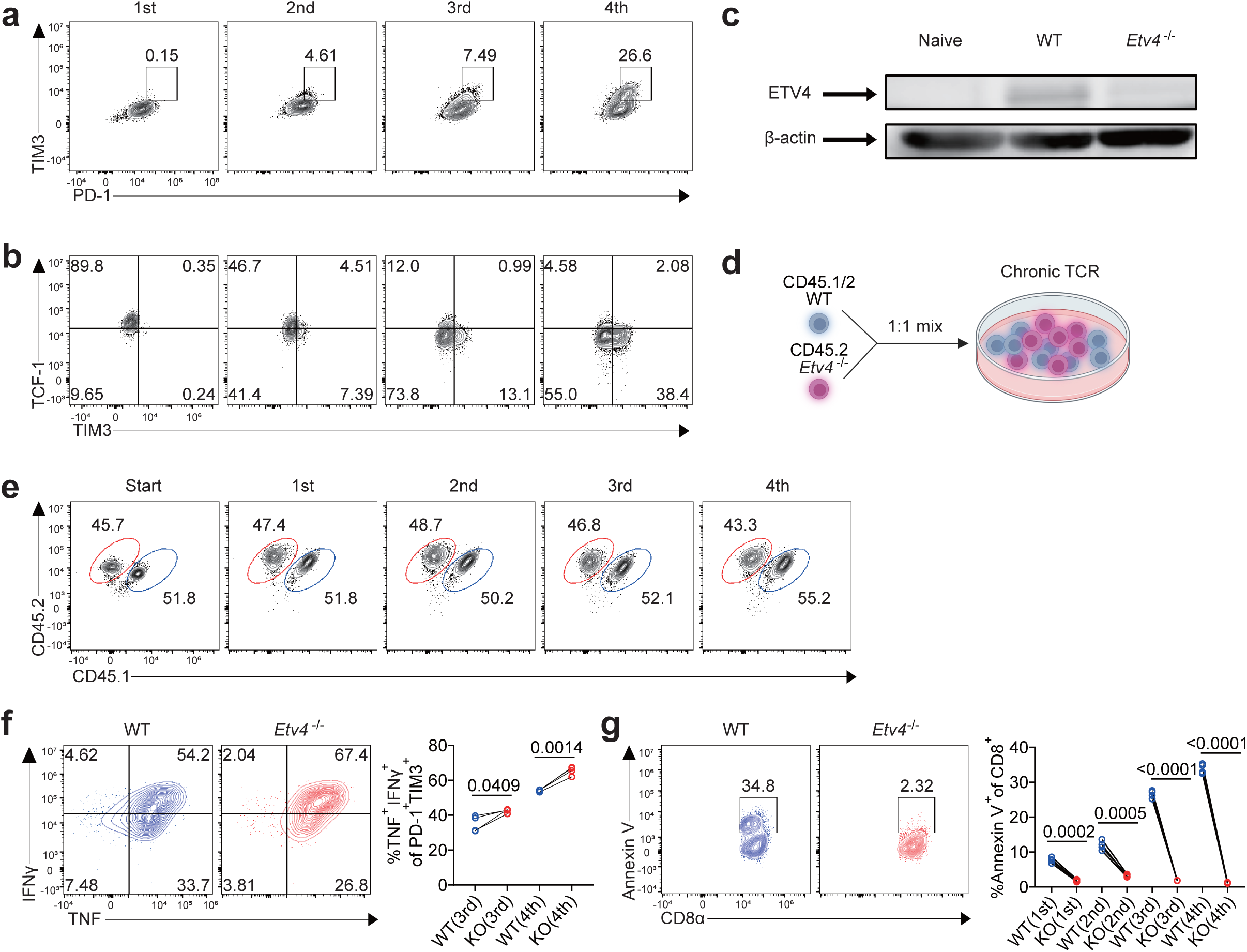
*In vitro* T cell exhaustion and co-culture of WT and *Etv4* KO CD8⁺ T cells. **a,** Representative flow cytometry plots of PD-1^hi^TIM3^+^ CD8^+^ T cells under *in vitro* chronic TCR stimulation. **b,** Representative flow cytometry plots of Tpex (TCF-1^+^TIM3^-^) or Tex cells (TCF-1^-^TIM3^+^) under chronic TCR stimulation. **c,** Western blot analysis of ETV4 levels in WT and *Etv4* KO CD8⁺ T cells after the fourth round of TCR stimulation, with naïve CD8⁺ T cells as negative control. **d,** Experimental outline of *in vitro* chronic TCR stimulation in co-culture of CD45.1/2 WT and CD45.2 *Etv4* KO CD8^+^ T cells. **e-f,** Representative flow cytometry plots and quantification of co-cultured CD45.1/2 WT (blue) and CD45.2 *Etv4* KO (red) (n=4) CD8^+^ T cells under *in vitro* chronic TCR stimulation: frequencies of WT and *Etv4* KO CD8^+^ T cells **(e)**, TNF^+^IFNγ^+^ frequencies within PD-1^hi^TIM3^+^ CD8^+^ cells at the third and fourth activation **(f)**, and Annexin V^+^ frequencies within CD8^+^ T cells **(g)**. The data represent three independent experiments for **a-c and e-g**. Error bars indicate mean ± s.e.m. Statistical significance was determined using paired two-tailed Student’s t-tests for **f-g**. Panel **d** was created using <u>BioRender.com</u>

**Extended Data Figure 2.**
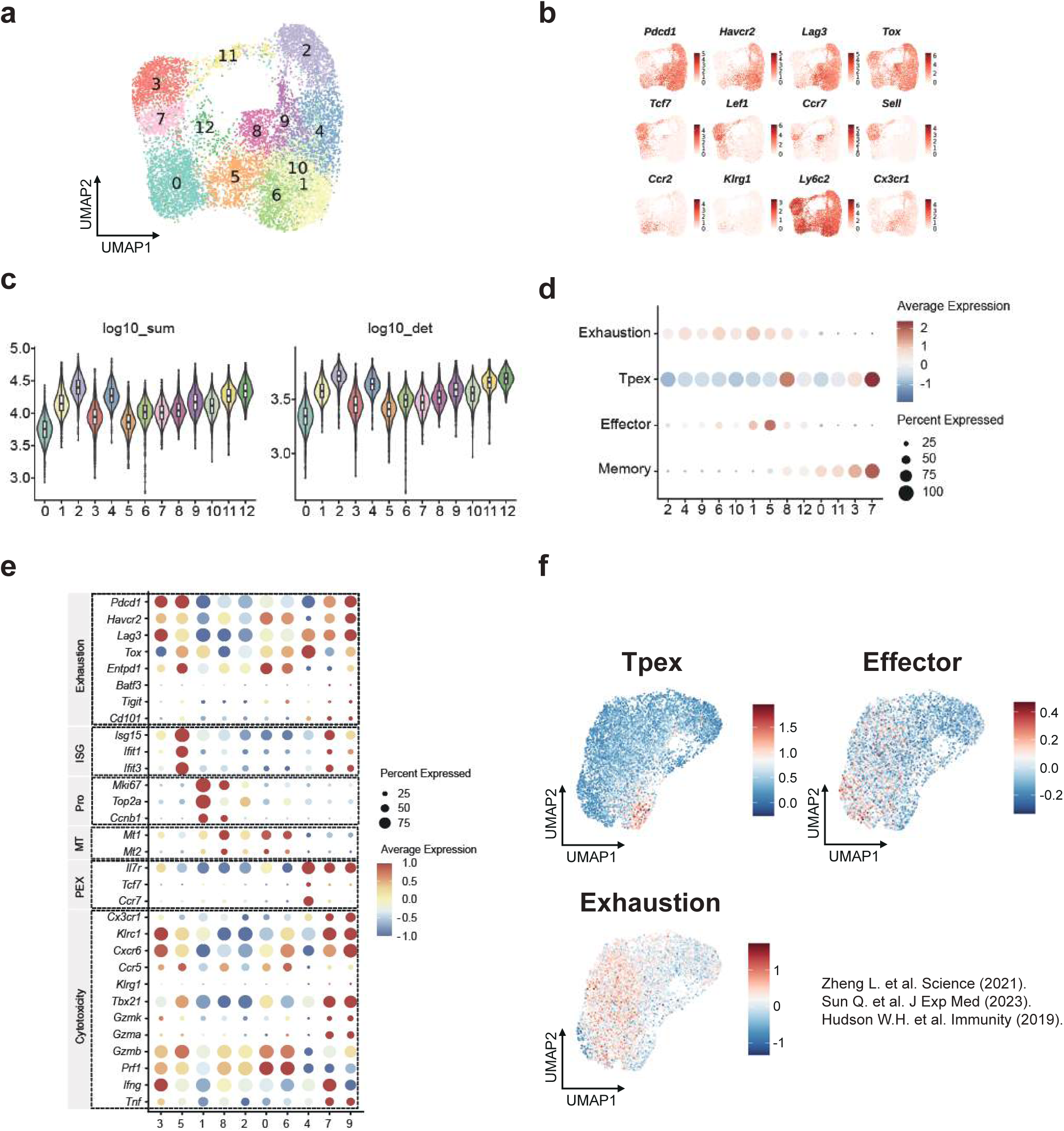
Validation of scRNA-seq data from MC38 tumor-infiltrating WT and *Etv4* KO CD8^+^ T cells. **a,** UMAP plot of CD8^+^ T cells from WT and *Etv4* KO mice. **b,** Feature plots of expression of canonical CD8^+^ T cell subtype marker genes. **c,** Violin plot of log_10_- scaled total number of unique molecular identifier (left) and log_10_-scaled number of detected genes (right) in each cluster. **d,** Dot plots of exhaustion-, Tpex-, effector-, and memory-associated gene signature scores in each cluster^28, 50, 78^. **e,** Dot plots of various genes in *Pdcd1*^+^ CD8⁺ T cells. **f,** UMAP visualization of CD8⁺ T cells from MC38 tumors showing effector-, exhaustion-, and Tpex-associated gene signature scores^28, 50, 78^. Color intensity reflects the relative enrichment of each signature per cell.

**Extended Data Figure 3.**
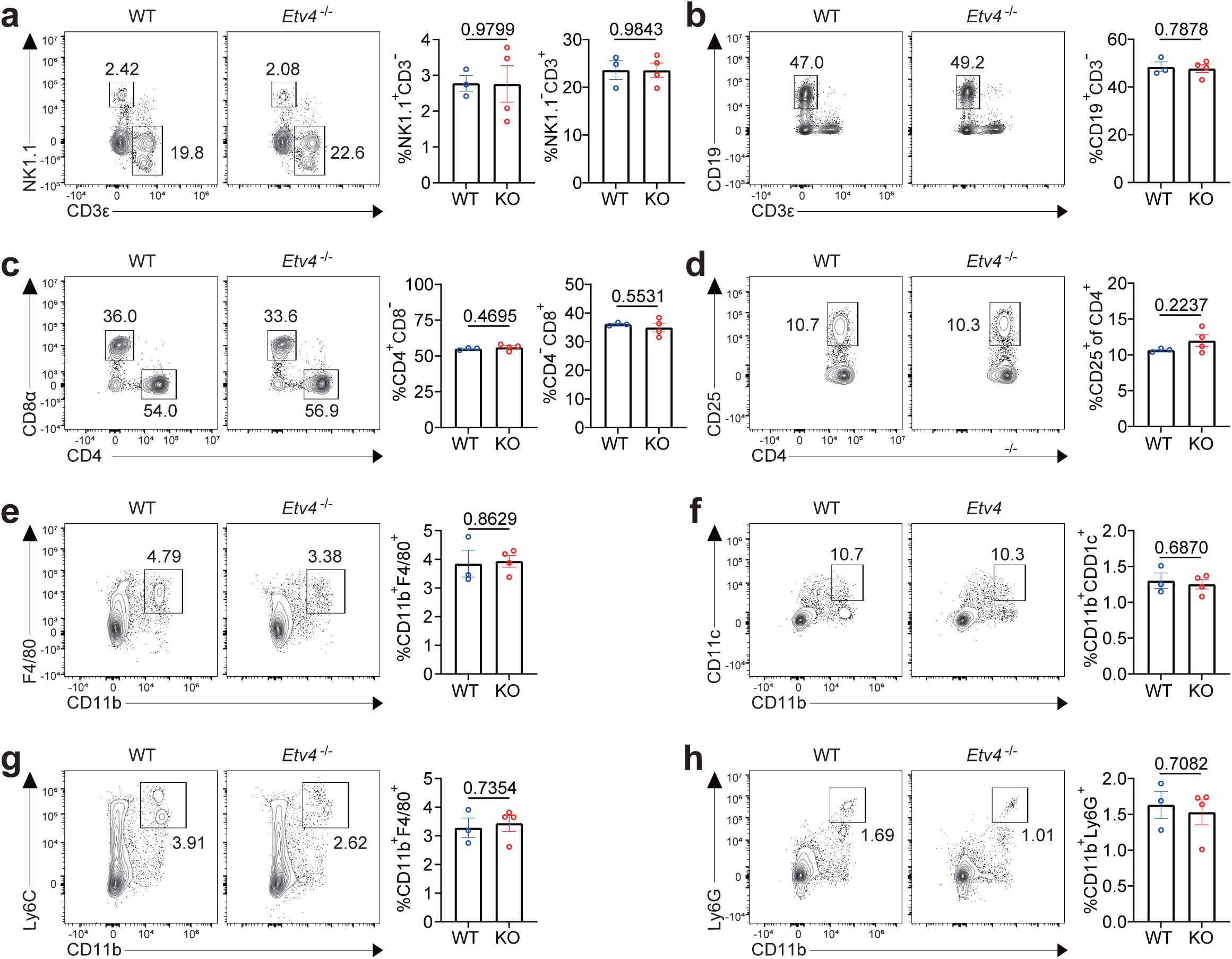
*Etv4* KO mice show no change in global immune cell populations at steady state. **a-h,** Representative flow cytometry plots and quantification of various immune cells in spleens of naïve WT (n=3) and *Etv4* KO (n=4) mice: natural killer cells (NK cells) (NK1.1^+^CD3^-^) and T cells (NK1.1^-^CD3^+^) **(a)**, B cells (CD19^+^CD3^-^) **(b),** CD4^+^ T cells and CD8^+^ T cells **(c),** CD25^+^CD4^+^ T cells **(d),** macrophage (CD11b^+^F4/80^+^) **(e)**, conventional dendritic cells (CD11b^+^CD11c^+^) **(f),** monocytes (CD11b^+^Ly6c^+^) **(g),** and neutrophils (CD11b^+^Ly6G^+^) **(h)**. The data represent three independent experiments for **a-h**. Error bars indicate mean ± s.e.m. Statistical significance was determined using unpaired two-tailed Student’s t-tests for **a-h**.

**Extended Data Figure 4.**
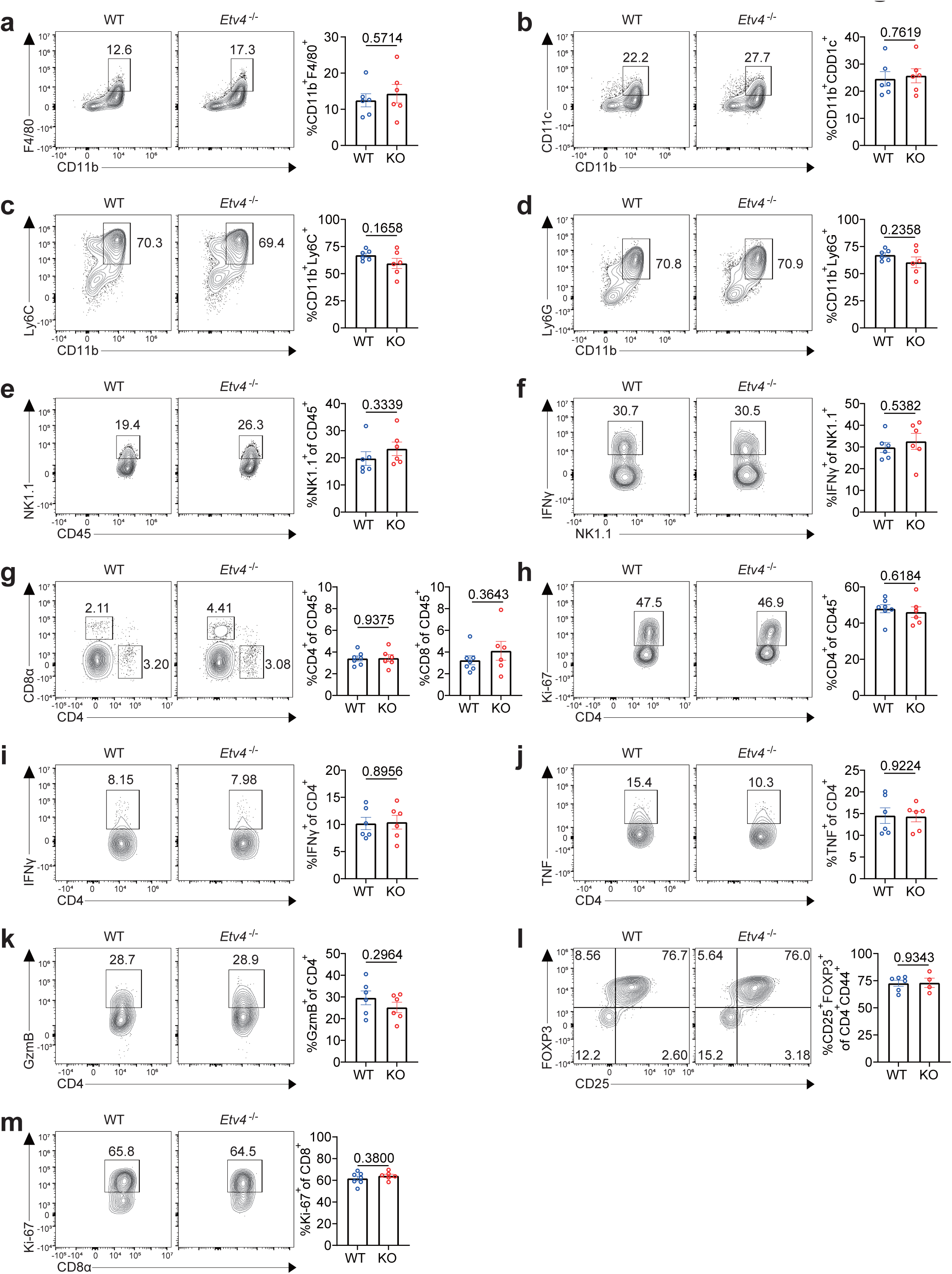
*Etv4* deficiency does not affect intratumoral myeloid cells, NK cells, and CD4^+^ T cells. **a-f,** Representative flow cytometry plots and quantification of various immune cells infiltrating MC38 tumors in WT (n=6) and *Etv4* KO (n=6) mice: macrophage (CD11b^+^F4/80^+^) **(a)**, dendritic cells (CD11b^+^CD11c^+^) **(b),** monocytes (CD11b^+^Ly6C^+^) **(c),** neutrophils (CD11b^+^Ly6G^+^) **(d),** NK cells (NK1.1^+^) **(e)**, and IFNγ^+^ NK cells, restimulated with PMA and ionomycin **(f). g-h,** Representative flow cytometry plots and quantification of CD4^+^ and CD8^+^ T cells infiltrating MC38 tumors in WT (n=7) and *Etv4* KO (n=6) mice: CD4^+^ T cells and CD8^+^ T cells **(g),** and Ki-67^+^ CD4^+^ T cells **(h). i-k,** Representative flow cytometry plots and quantification of CD4^+^ T cells infiltrating MC38 tumors in WT (n=6) and *Etv4* KO (n=6) mice, restimulated with PMA and ionomycin: IFNγ^+^ cells **(i),** TNF^+^ cells **(j),** and granzyme B^+^ cells **(k). l,** Representative flow cytometry plots and quantification of CD25^+^FOXP3^+^ regulatory CD4^+^ T cells infiltrating MC38 tumors in WT (n=6) and *Etv4* KO (n=4) mice. **m,** Representative flow cytometry plots and quantification of Ki-67^+^ CD8^+^ T cells infiltrating MC38 tumors in WT (n=7) and *Etv4* KO (n=6) mice. The data represent three independent experiments for **a-m**. Error bars indicate mean ± s.e.m. Statistical significance was determined using unpaired two-tailed Student’s t-tests for **a-m**.

**Extended Data Figure 5.**
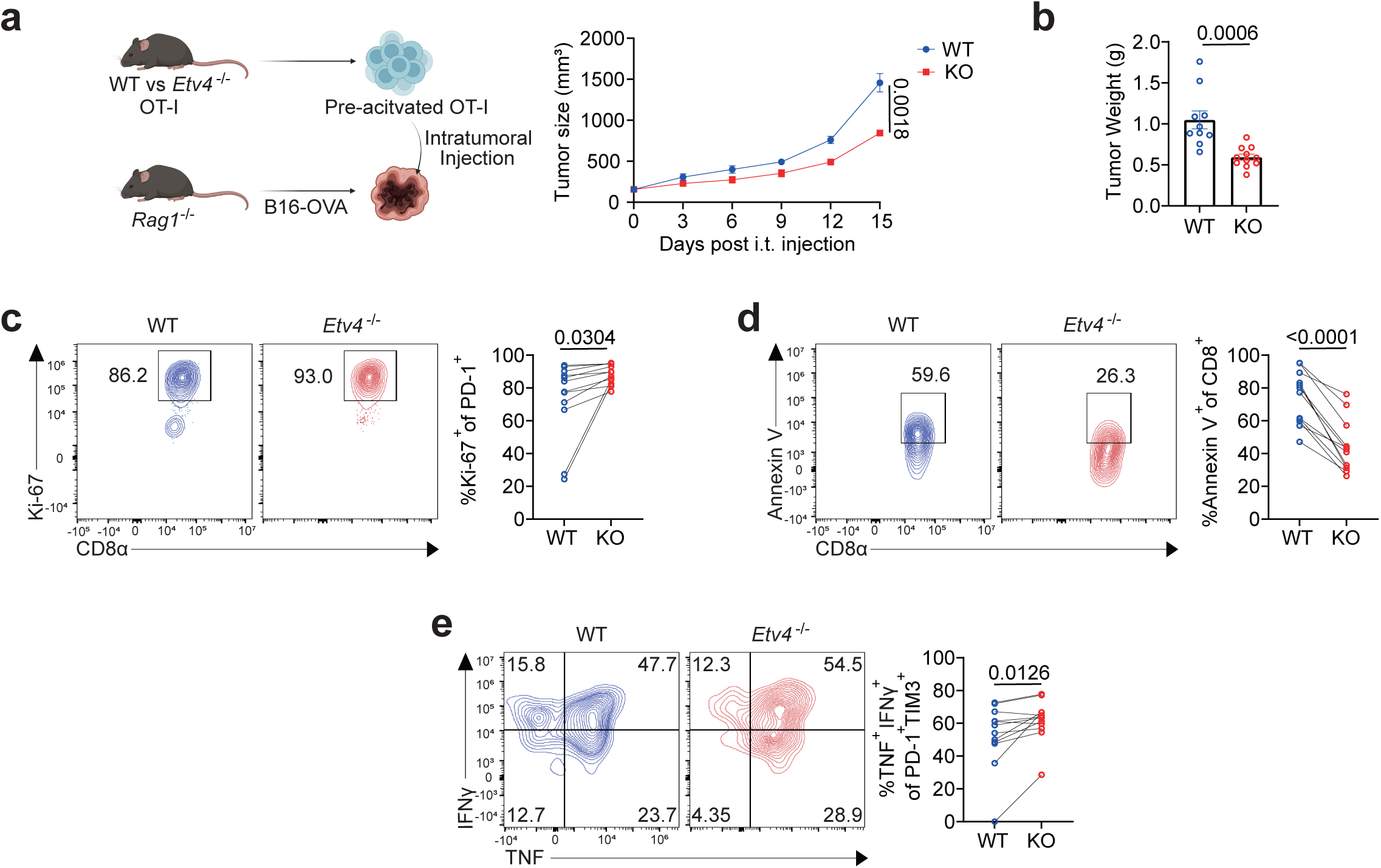
Assessment of the intrinsic contribution of ETV4 to CD8⁺ T cell-mediated tumor control. **a**, Experimental outline of intratumoral adoptive transfer of WT or *Etv4* KO OT-I cells into B16-OVA tumor-bearing *Rag1* KO mice. Tumor growth curves in B16-OVA tumor-bearing *Rag1* KO mice intratumorally adoptively transferred with WT OT-I (n=10) or *Etv4* KO OT-I (n=11) cells. **b,** Tumor weight of B16-OVA tumor-bearing *Rag1* KO mice following intratumoral adoptive transfer of WT OT-I (n=10) and *Etv4* KO OT-I (n=11) cells. **c-e,** Representative flow cytometry plots and quantification of OT-I cells in B16-OVA tumor-bearing *Rag1* KO mice after intratumoral co-adoptive transfer of CD45.1 WT (blue) and CD45.2 *Etv4* KO OT-I cells (red) (n=12): Ki-67⁺ cells **(c)**, Annexin V^+^ cells **(d),** and TNF⁺IFNγ⁺ frequencies within PD-1^hi^TIM3^+^ CD8^+^ cells **(e).** The data represent three independent experiments for **a-e**. Error bars indicate mean ± s.e.m. Statistical significance was determined using a two-way ANOVA with Tukey’s multiple comparison for **a**, unpaired two-tailed Student’s t-tests for **b,** and paired for **c-e.** Data were pooled from two (**a-b**) independent experiments. Panel **a** was created using <u>BioRender.com</u>

**Extended Data Figure 6.**
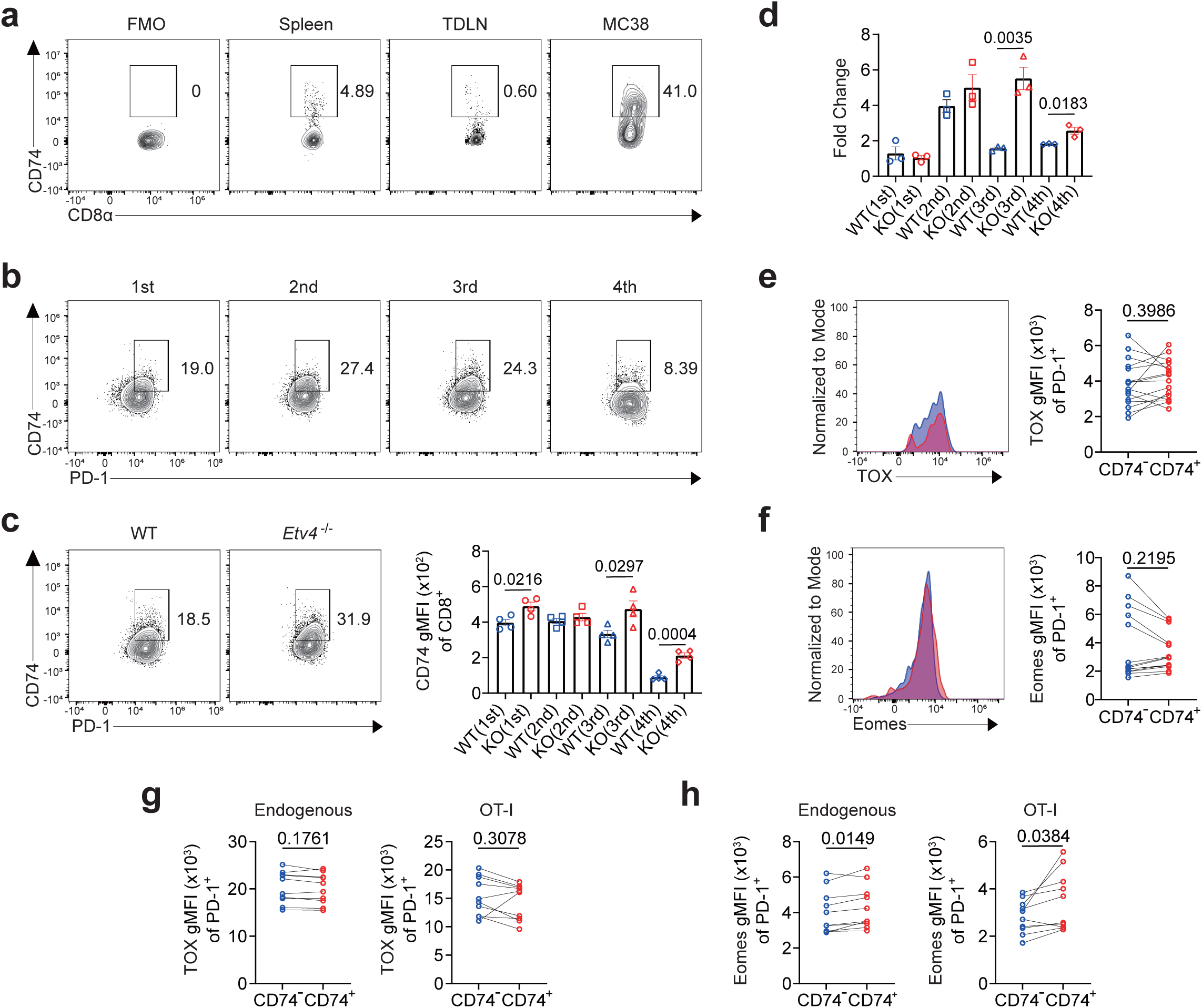
Phenotypic and transcriptional characterization of CD74⁺ CD8⁺ T cells. **a,** Representative flow cytometry plots of CD74^+^ CD8^+^ T cells in spleen, tumor-draining lymph nodes, and MC38 tumors. **b,** Representative flow cytometry plots of CD74^+^ CD8^+^ T cells under *in vitro* chronic TCR stimulation. **c,** Representative flow cytometry plots at the third activation and gMFI quantification of CD74 surface expression in CD8^+^ T cells from WT (n=4) and *Etv4* KO (n=4). **d,** RT-qPCR analysis for *Cd74* expression levels in WT (n=3) and *Etv4* KO (n=3) CD8⁺ T cells under *in vitro* chronic TCR stimulation. **e-f,** Representative histogram and gMFI quantification of intracellular transcription factor expression in CD74^-^ (blue) versus CD74^+^ (red) PD-1^+^ CD8^+^ T cells from MC38 tumor-bearing mice: TOX (n=16) **(e)**, and Eomes (n=14) **(f). g-h,** gMFI quantification of intracellular transcription factor expression in CD74⁻ versus CD74⁺PD-1⁺ cells from CD45.2 endogenous CD8^+^ and CD45.1/2 OT-I cells (intratumorally adoptive transfer) infiltrating B16-OVA tumor-bearing mice (n=10): TOX **(g)** and Eomes **(h).** The data represent three independent experiments for **a-h**. Error bars indicate mean ± s.e.m. Statistical significance was determined using unpaired two- tailed Student’s t-tests for **c-d,** and paired for **e-h**. Data were pooled from 2 (**e-f**) independent experiments.

**Extended Data Figure 7.**
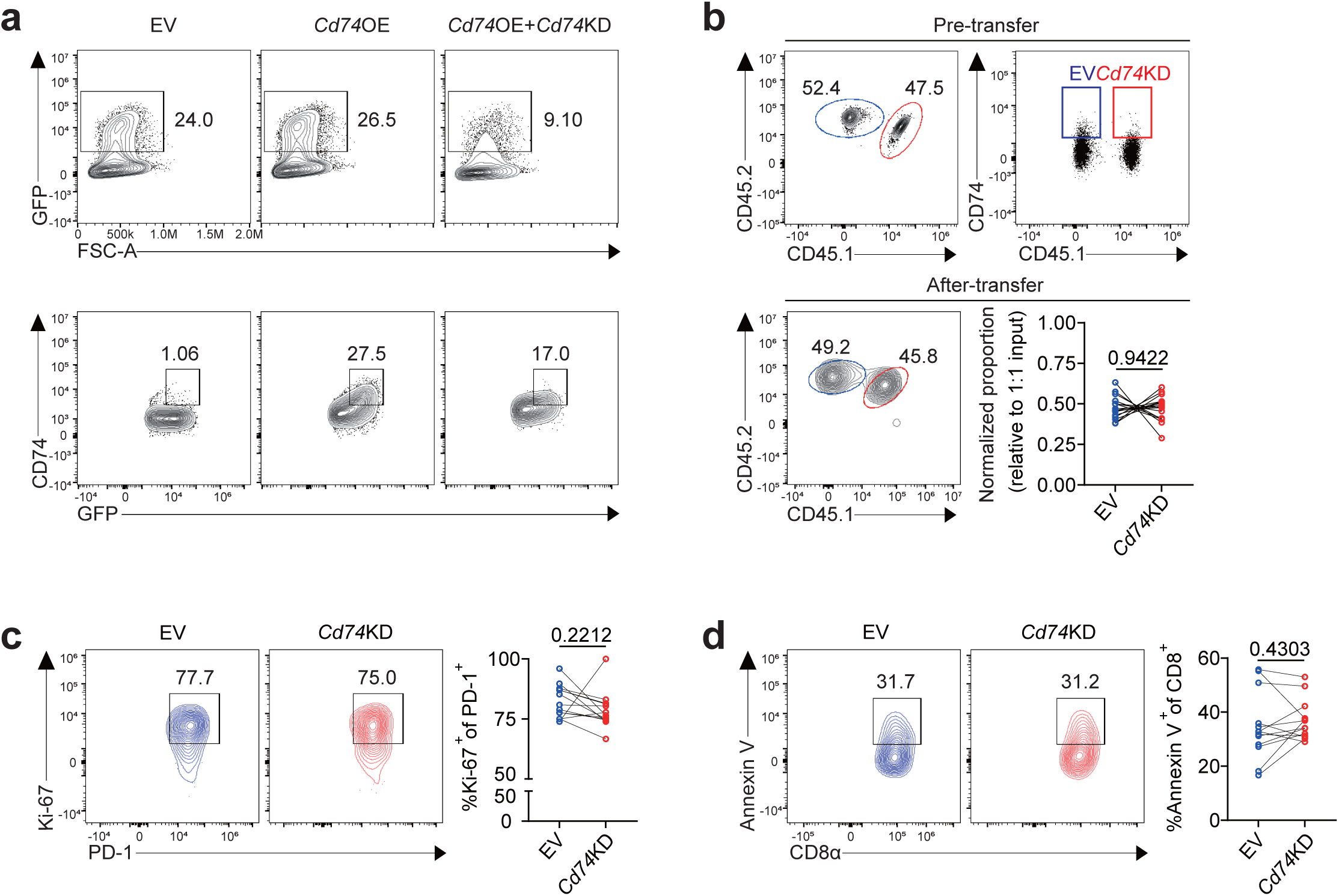
CD74 loss has no effect on CD8⁺ T cell proliferation or apoptosis. **a,** Representative flow cytometry plots of CD74^+^GFP⁺ HEK293T cells transfected with EV, *Cd74* OE, or *Cd74* OE + *Cd74* KD. **b-d,** Representative flow cytometry plots and quantification of OT-I cells in B16-OVA tumor-bearing CD45.1 immunocompetent mice after intratumoral co-adoptive transfer of CD45.2 EV (blue) and CD45.1/2 *Cd74* KD OT-I T cells (red) (n=13): validation of 1:1 co-adoptive pre-transfer and normalized proportion after transfer **(b)**, Ki-67^+^ cells **(c),** and Annexin V^+^ cells **(d).** The data represent one experiment for **a,** and three independent experiments for **b-d**. Error bars indicate mean ± s.e.m. Statistical significance was determined using paired two-tailed Student’s t-tests for **b-d**.

**Extended Data Figure 8.**
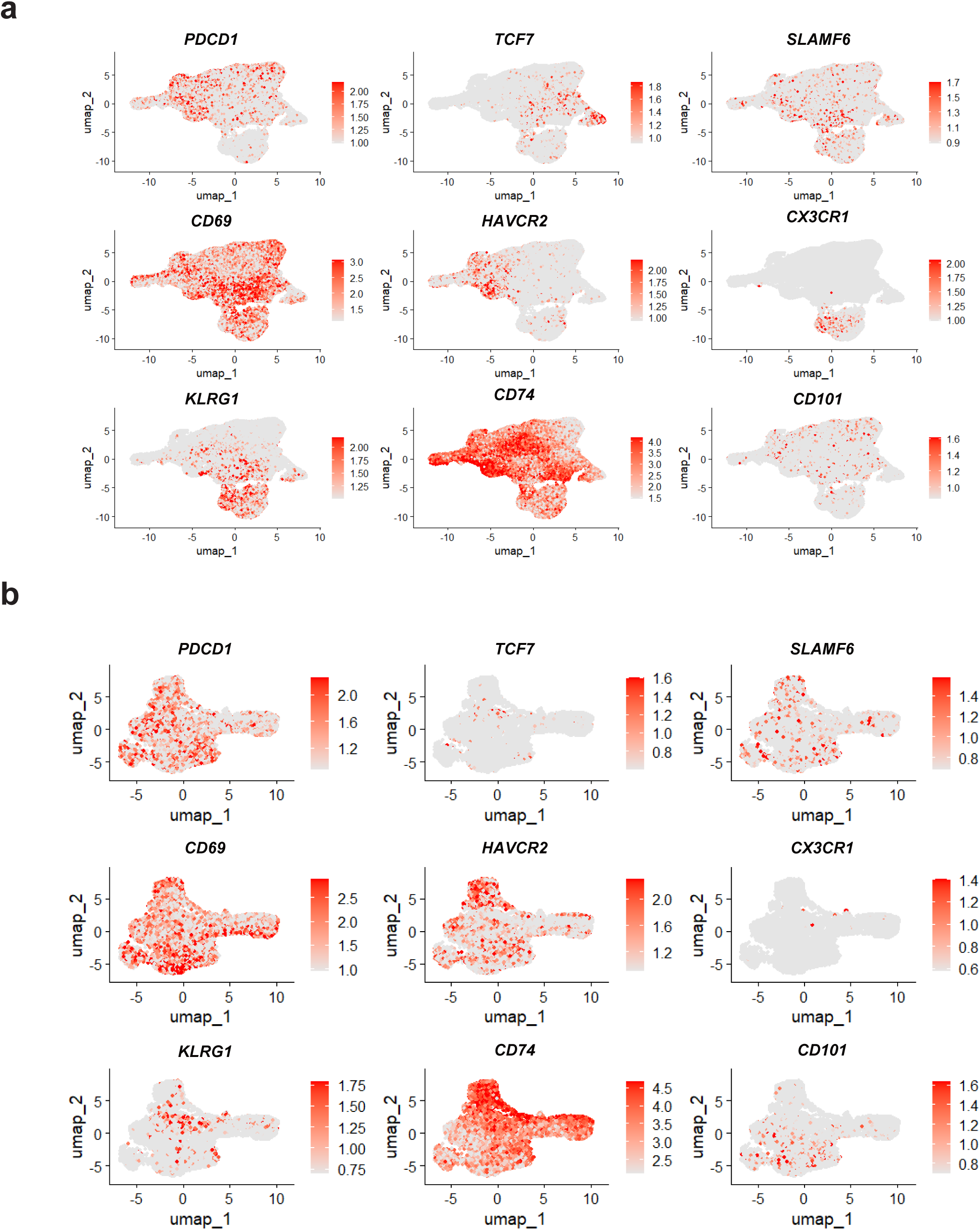
Exhaustion-associated gene expression patterns in human pan-cancer. **a,** Feature plots showing exhaustion- associated gene expression (red) across initial UMAP clusters of CD8⁺ T cells^50^. **b,** Feature plots showing exhaustion-associated gene expression (red) across subclustered UMAP clusters of *PDCD1*^+^*HAVCR2*^+^ CD8*^+^* T cells.

**Extended Data Figure 9.**
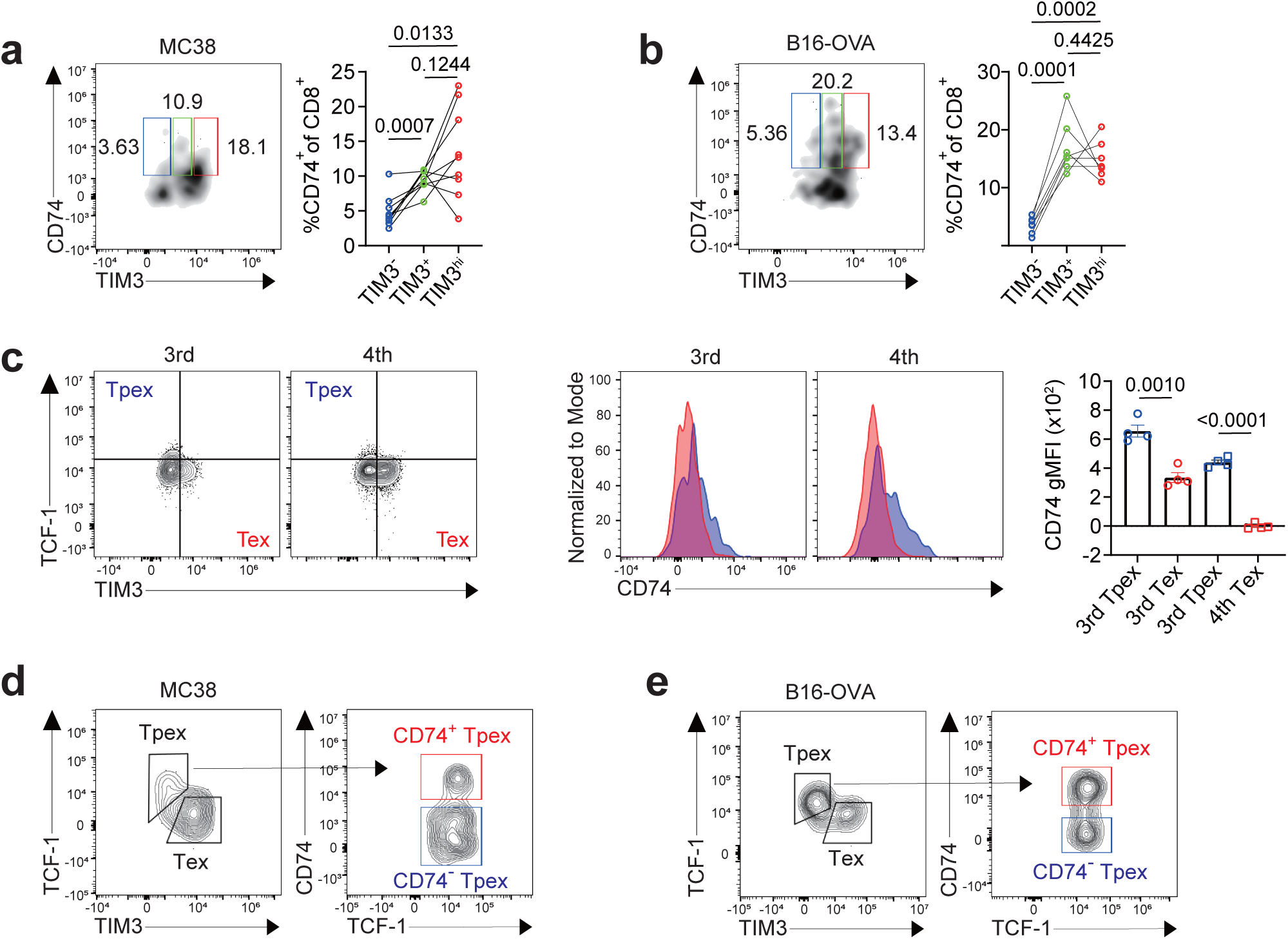
CD74 expression is regulated by environmental cues and distinguishes Tpex subsets that may harbor divergent potential. **a-b,** Representative flow cytometry plots and quantification of CD74^+^ cells from CD8^+^ T cells infiltrating tumors in WT mice: MC38 (n=9) **(a)**, and B16-OVA (n=7) **(b). c,** Representative flow cytometry plots, histograms, and gMFI quantification of CD74 surface expression from CD8^+^ T cells at the third and fourth activation in the *in vitro* T cell exhaustion model (n=4). **d-e,** Representative flow cytometry plots of CD74^+^ and CD74^-^ Tpex (TCF-1^+^TIM3^-^) cells infiltrating tumors in WT mice: MC38 **(d)** and B16-OVA **(e).** The data represent three independent experiments for **a-e**. Error bars indicate mean ± s.e.m. Statistical significance was determined using paired two-tailed Student’s t-tests for **a-b** and unpaired for **c**.

## REFERENCES

1. Dolina, J.S., Van Braeckel-Budimir, N., Thomas, G.D. & Salek-Ardakani, S. CD8+ T cell exhaustion in cancer. Frontiers in immunology 12, 715234 (2021).

2. Lichterfeld, M. et al. Loss of HIV-1–specific CD8+ T cell proliferation after acute HIV-1 infection and restoration by vaccine-induced HIV-1–specific CD4+ T cells. The Journal of experimental medicine 200, 701–712 (2004).

3. Guan, Q. et al. Strategies to reinvigorate exhausted CD8+ T cells in tumor microenvironment. Frontiers in Immunology 14, 1204363 (2023).

4. Miller, B.C. et al. Subsets of exhausted CD8+ T cells differentially mediate tumor control and respond to checkpoint blockade. Nature immunology 20, 326–336 (2019).

5. Paley, M.A. et al. Progenitor and terminal subsets of CD8+ T cells cooperate to contain chronic viral infection. Science 338, 1220–1225 (2012).

6. Jiang, Y., Li, Y. & Zhu, B. T-cell exhaustion in the tumor microenvironment. Cell death & disease 6, e1792–e1792 (2015).

7. Guo, Y. et al. Metabolic reprogramming of terminally exhausted CD8+ T cells by IL-10 enhances anti-tumor immunity. Nature immunology 22, 746–756 (2021).

8. Gill, A.L. et al. PD-1 blockade increases the self-renewal of stem-like CD8 T cells to compensate for their accelerated differentiation into effectors. Science immunology 8, eadg0539 (2023).

9. Hashimoto, M. et al. PD-1 combination therapy with IL-2 modifies CD8+ T cell exhaustion program. Nature 610, 173–181 (2022).

10. Khan, O. et al. TOX transcriptionally and epigenetically programs CD8+ T cell exhaustion. Nature 571, 211–218 (2019).

11. Yoshikawa, T. et al. Genetic ablation of PRDM1 in antitumor T cells enhances therapeutic efficacy of adoptive immunotherapy. *Blood*, The Journal of the American Society of Hematology 139, 2156–2172 (2022).

12. Scharping, N.E. et al. Mitochondrial stress induced by continuous stimulation under hypoxia rapidly drives T cell exhaustion. Nature immunology 22, 205–215 (2021).

13. Kao, C. et al. T-bet represses expression of PD-1 and sustains virus-specific CD8 T cell responses during chronic infection. Nature immunology 12, 663 (2011).

14. Li, J., He, Y., Hao, J., Ni, L. & Dong, C. High levels of Eomes promote exhaustion of anti-tumor CD8+ T cells. Frontiers in immunology 9, 2981 (2018).

15. Chen, Y. et al. BATF regulates progenitor to cytolytic effector CD8+ T cell transition during chronic viral infection. Nature immunology 22, 996–1007 (2021).

16. Huang, Y.J. et al. Continuous expression of TOX safeguards exhausted CD8 T cell epigenetic fate. Science Immunology 10, eado3032 (2025).

17. Damo, M. et al. PD-1 maintains CD8 T cell tolerance towards cutaneous neoantigens. Nature 619, 151–159 (2023).

18. Weiss, S.A. et al. Epigenetic tuning of PD-1 expression improves exhausted T cell function and viral control. Nature Immunology 25, 1871–1883 (2024).

19. Dumortier, M. et al. ETV4 transcription factor and MMP13 metalloprotease are interplaying actors of breast tumorigenesis. Breast Cancer Research 20, 1–18 (2018).

20. Aytes, A. et al. ETV4 promotes metastasis in response to activation of PI3-kinase and Ras signaling in a mouse model of advanced prostate cancer. Proceedings of the National Academy of Sciences 110, E3506–E3515 (2013).

21. Zhu, T. et al. ETV4 promotes breast cancer cell stemness by activating glycolysis and CXCR4-mediated sonic Hedgehog signaling. Cell death discovery 7, 126 (2021).

22. Kim, E. et al. Capicua suppresses hepatocellular carcinoma progression by controlling the ETV4–MMP1 axis. Hepatology 67, 2287–2301 (2018).

23. Lee, J.-S. et al. Capicua suppresses colorectal cancer progression via repression of ETV4 expression. Cancer Cell International 20, 42 (2020).

24. Zhao, Y. et al. Single-cell transcriptomics of immune cells reveal diversity and exhaustion signatures in non-small-cell lung cancer. Frontiers in Immunology 13, 854724 (2022).

25. Ma, S. et al. Nutrient-driven histone code determines exhausted CD8+ T cell fates. Science 387, eadj3020 (2024).

26. Humblin, E. et al. The costimulatory molecule ICOS limits memory-like properties and function of exhausted PD-1+ CD8+ T cells. Immunity 58, 1966–1983. e1910 (2025).

27. Doedens, A.L. et al. Hypoxia-inducible factors enhance the effector responses of CD8+ T cells to persistent antigen. Nature immunology 14, 1173–1182 (2013).

28. Hudson, W.H. et al. Proliferating transitory T cells with an effector-like transcriptional signature emerge from PD-1+ stem-like CD8+ T cells during chronic infection. Immunity 51, 1043–1058. e1044 (2019).

29. Daniel, B. et al. Divergent clonal differentiation trajectories of T cell exhaustion. Nature immunology 23, 1614–1627 (2022).

30. Minnie, S.A. et al. TIM-3+ CD8 T cells with a terminally exhausted phenotype retain functional capacity in hematological malignancies. Science immunology 9, eadg1094 (2024).

31. Pauken, K.E. et al. Single-cell analyses identify circulating anti-tumor CD8 T cells and markers for their enrichment. Journal of Experimental Medicine 218, e20200920 (2021).

32. Vignali, P.D. et al. Hypoxia drives CD39-dependent suppressor function in exhausted T cells to limit antitumor immunity. Nature immunology 24, 267–279 (2023).

33. Siddiqui, I. et al. Intratumoral Tcf1+ PD-1+ CD8+ T cells with stem-like properties promote tumor control in response to vaccination and checkpoint blockade immunotherapy. Immunity 50, 195–211. e110 (2019).

34. Su, H., Na, N., Zhang, X. & Zhao, Y. The biological function and significance of CD74 in immune diseases. Inflammation Research 66, 209–216 (2017).

35. Strijker, J.G. et al. Blocking MIF secretion enhances CAR T-cell efficacy against neuroblastoma. European Journal of Cancer 218, 115263 (2025).

36. Bonnin, E. et al. CD74 supports accumulation and function of regulatory T cells in tumors. Nature Communications 15, 3749 (2024).

37. Fan, H. et al. Macrophage migration inhibitory factor and CD74 regulate macrophage chemotactic responses via MAPK and Rho GTPase. The Journal of Immunology 186, 4915–4924 (2011).

38. Pellegrino, B. et al. CD74 promotes the formation of an immunosuppressive tumor microenvironment in triple-negative breast cancer in mice by inducing the expansion of tolerogenic dendritic cells and regulatory B cells. PLoS Biology 22, e3002905 (2024).

39. Figueiredo, C.R. et al. Blockade of MIF–CD74 signalling on macrophages and dendritic cells restores the antitumour immune response against metastatic melanoma. Frontiers in immunology 9, 1132 (2018).

40. Beltra, J.-C. et al. Developmental relationships of four exhausted CD8+ T cell subsets reveals underlying transcriptional and epigenetic landscape control mechanisms. Immunity 52, 825–841. e828 (2020).

41. Westmeier, J. et al. Macrophage migration inhibitory factor receptor CD74 expression is associated with expansion and differentiation of effector T cells in COVID-19 patients. Frontiers in Immunology 14, 1236374 (2023).

42. Tanese, K. et al. Cell surface CD74–MIF interactions drive melanoma survival in response to interferon-γ. Journal of investigative dermatology 135, 2775–2784 (2015).

43. Intlekofer, A.M. et al. Effector and memory CD8+ T cell fate coupled by T-bet and eomesodermin. Nature immunology 6, 1236–1244 (2005).

44. Szabo, S.J. et al. A novel transcription factor, T-bet, directs Th1 lineage commitment. Cell 100, 655–669 (2000).

45. Oh, S. & Hwang, E.S. The role of protein modifications of T-bet in cytokine production and differentiation of T helper cells. Journal of immunology research 2014, 589672 (2014).

46. Hatton, R.D. et al. A distal conserved sequence element controls Ifng gene expression by T cells and NK cells. Immunity 25, 717–729 (2006).

47. Sullivan, B.M., Juedes, A., Szabo, S.J., Von Herrath, M. & Glimcher, L.H. Antigen-driven effector CD8 T cell function regulated by T-bet. Proceedings of the National Academy of Sciences 100, 15818–15823 (2003).

48. Mayer, K.D. et al. Cutting edge: T-bet and IL-27R are critical for in vivo IFN-γ production by CD8 T cells during infection. The Journal of Immunology 180, 693–697 (2008).

49. Yang, Y., Ochando, J.C., Bromberg, J.S. & Ding, Y. Identification of a distant T-bet enhancer responsive to IL-12/Stat4 and IFNγ/Stat1 signals. *Blood*, The Journal of the American Society of Hematology 110, 2494–2500 (2007).

50. Zheng, L. et al. Pan-cancer single-cell landscape of tumor-infiltrating T cells. Science 374, abe6474 (2021).

51. Hough, K.P., Chisolm, D.A. & Weinmann, A.S. Transcriptional regulation of T cell metabolism. Molecular immunology 68, 520–526 (2015).

52. Matsui, M., Moriya, O., Yoshimoto, T. & Akatsuka, T. T-bet is required for protection against vaccinia virus infection. Journal of virology 79, 12798–12806 (2005).

53. Rosas, L.E. et al. Cutting edge: STAT1 and T-bet play distinct roles in determining outcome of visceral leishmaniasis caused by Leishmania donovani. The Journal of Immunology 177, 22–25 (2006).

54. Juedes, A.E., Rodrigo, E., Togher, L., Glimcher, L.H. & von Herrath, M.G. T-bet controls autoaggressive CD8 lymphocyte responses in type 1 diabetes. The Journal of experimental medicine 199, 1153–1162 (2004).

55. Bertrand, F. et al. Blocking Tumor Necrosis Factor α Enhances CD8 T-cell–dependent immunity in experimental melanoma. Cancer research 75, 2619–2628 (2015).

56. Liu, Y. et al. IL-2 regulates tumor-reactive CD8+ T cell exhaustion by activating the aryl hydrocarbon receptor. Nature immunology 22, 358–369 (2021).

57. Huseni, M.A., et al. CD8+ T cell-intrinsic IL-6 signaling promotes resistance to anti-PD-L1 immunotherapy. Cell Reports Medicine 4 (2023).

58. Hu, Y. et al. TGF-β regulates the stem-like state of PD-1+ TCF-1+ virus-specific CD8 T cells during chronic infection. Journal of Experimental Medicine 219, e20211574 (2022).

59. Chen, W. et al. Chronic type I interferon signaling promotes lipid-peroxidation-driven terminal CD8+ T cell exhaustion and curtails anti-PD-1 efficacy. Cell reports 41 (2022).

60. Sumida, T.S. et al. Type I interferon transcriptional network regulates expression of coinhibitory receptors in human T cells. Nature immunology 23, 632–642 (2022).

61. Hwang, S.-M. et al. Transgelin 2 guards T cell lipid metabolism and antitumour function. Nature 635, 1010–1018 (2024).

62. Faust Akl, C., et al. Glioblastoma-instructed astrocytes suppress tumour-specific T cell immunity. Nature, 1–11 (2025).

63. Noe, J.T. & Mitchell, R.A. MIF-dependent control of tumor immunity. Frontiers in immunology 11, 609948 (2020).

64. Niborski, L.L. et al. CD8+ T cell responsiveness to anti-PD-1 is epigenetically regulated by Suv39h1 in melanomas. Nature communications 13, 3739 (2022).

65. Gil-Yarom, N. et al. CD74 is a novel transcription regulator. Proceedings of the National Academy of Sciences 114, 562–567 (2017).

66. Ma, C. et al. Therapeutic modulation of APP-CD74 axis can activate phagocytosis of TAMs in GBM. Biochimica et Biophysica Acta (BBA)-Molecular Basis of Disease 1870, 167449 (2024).

67. Song, Y. & Lee, Y. Brief guide to flow cytometry. Molecules and Cells 47, 100129 (2024).

68. Bong, D., Sohn, J. & Lee, S.-J.V. Brief guide to RT-qPCR. Molecules and Cells 47, 100141 (2024).

69. Badia, I.M.P., et al. decoupleR: ensemble of computational methods to infer biological activities from omics data. Bioinform Adv 2, vbac016 (2022).

70. Korotkevich, G. et al. Fast gene set enrichment analysis. bioRxiv, 060012 (2021).

71. Dolgalev, I. msigdbr: MSigDB Gene Sets for Multiple Organisms in a Tidy Data Format. 2025.

72. Zheng, G.X. et al. Massively parallel digital transcriptional profiling of single cells. Nat Commun 8, 14049 (2017).

73. Lun, A.T.L. et al. EmptyDrops: distinguishing cells from empty droplets in droplet-based single-cell RNA sequencing data. Genome Biol 20, 63 (2019).

74. McCarthy, D.J., Campbell, K.R., Lun, A.T. & Wills, Q.F. Scater: pre-processing, quality control, normalization and visualization of single-cell RNA-seq data in R. Bioinformatics 33, 1179–1186 (2017).

75. Lun, A.T., Bach, K. & Marioni, J.C. Pooling across cells to normalize single-cell RNA sequencing data with many zero counts. Genome Biol 17, 75 (2016).

76. Hao, Y. et al. Dictionary learning for integrative, multimodal and scalable single-cell analysis. Nat Biotechnol 42, 293–304 (2024).

77. Germain, P.L., Lun, A., Garcia Meixide, C., Macnair, W. & Robinson, M.D. Doublet identification in single-cell sequencing data using scDblFinder. F1000Res 10, 979 (2021).

78. Sun, Q. et al. STAT3 regulates CD8+ T cell differentiation and functions in cancer and acute infection. Journal of Experimental Medicine 220, e20220686 (2023).

